# Synchrony-driven Assemblies Reliably Represent Complex Stimuli in Ferret Auditory Cortex

**DOI:** 10.1101/2025.10.17.683032

**Authors:** Luca Rakowski

## Abstract

Neuronal assemblies defined by coordination at high temporal resolution are thought to act as functional modules for information processing throughout the brain. Here, we develop a new method for identifying these assemblies from analytical tests of pairwise synchrony, benefiting from three rigorous criteria for assembly detection and sensitivity to rare but significant coordinated firing patterns. Following validation through simulations and in vivo recordings, we perform synchrony-based analysis on datasets from both primary (A1) and non-primary (PEG) ferret auditory cortex during passive listening to complex natural sounds. We show that synchrony-driven assemblies (SDAs) in both areas exhibit regular temporal dynamics, including the production of stereotypical spike sequences. In-keeping with their enhanced inter-neuronal reliability, SDAs are observed to outperform random assemblies in two versions of rate-based decoding. We extend these results to temporal decoding, and consistently observe optimal integration windows of 10-20ms. Thus, we suggest that SDAs are sufficient for the accurate representation of diverse sets of complex auditory stimuli, and that this increase in information depends entirely on coordination at very fine temporal scales.

## Introduction

Neuronal representations of sensory information are commonly studied on either the level of single units or whole populations. In many cases, single units are sufficient to convey highly multiplexed readouts of complex stimuli. In others, individual neurons remain unreliable and require interpretation through population responses. The duality of these two families of methods imposes a false dichotomy that ignores the interrogation of small networks. Recently, there has been significant interest in the study of these networks - termed assemblies (Buzsáki, 2010; Miehl et al., 2022) - and how their member neurons may be optimally chosen to isolate the greatest amount of coherent information in the fewest cells (Ince, Panzeri, and Kayser, 2013). In theory, assemblies may represent functional modules that provide an additional level of organization to cortical computational processes (Buzsáki, 2010; Papadimitriou et al., 2020).

Assembly analysis has not escaped the field of auditory neuroscience. Consistent correlation structures in auditory cortex neurons, for example, have been shown to reflect spatial clustering (Eggermont, 2006; C. A. Atencio and C. E. Schreiner, 2013). Similarly, the identification of repeating patterns of cortical activity during complex cognitive tasks is again indicative of assembly-like organization (Villa et al., 1998). More recently, an explicit search for assemblies in auditory cortex revealed that stable synchronous activity (<10ms) could not be entirely explained by spectro-temporal receptive fields (STRFs), suggesting that assemblies in this area may encode more complex properties of sound (See et al., 2018). This hypothesis, however, has yet to be tested.

To answer this question, we analyze data obtained from primary (A1) and non-primary (PEG) ferret auditory cortex (Pennington and Stephen V. David, 2022) during passive listening. First, we aim to characterize network dynamics through a neuron clustering protocol reliant on the detection of non-spurious pairwise coincident spiking activity. To achieve this, we build on spike dithering (Date, Bienenstock, and Geman, 1998) as a surrogate means of significance testing. The matrix Δ of significant pairwise coactivity scores *δ*_*i,j*_ is then used to search for putative assemblies *E*. This approach is validated in both simulated and ferret datasets. Finally, we aim to understand whether and how assemblies benefit from spike synchrony in different coding schemes. We therefore interrogate ferret assemblies in two forms of rate coding by comparing their performance with that of randomly generated assemblies. Finally, we quantify the ability of synchrony-driven assemblies to encode complex sounds with temporal codes.

## Methods

In this section, we describe a process for performing pairwise synchrony analysis, for significance testing, and for developing the matrix of pairwise results into a basis for assembly discovery. We then discuss simulation methodology and methods pertaining to rate code and temporal code analysis. All experiments presented herein are conducted in accordance with these methods. Data from ferret auditory cortex was obtained from Pennington and Stephen V. David, 2022. Error bars represent 95% confidence intervals unless otherwise stated.

### Pairwise Coactivity Scores (*δ*)

There exist many ways of computing a coactivity score *δ*_*i,j*_ between a pair of spike trains *i, j* (Pipa, Wheeler, et al., 2008). A common solution is to compute Pearson’s correlation coefficient from binned and z-scored spike trains (Peyrache et al., 2009; Lopesdos-Santos, Ribeiro, and Tort, 2013). However, binning can underestimate correlation due to artificial separation at bin margins, or overestimate synchrony when bins are too wide. Further, correlation analysis is sensitive to sparsity due to most principal cells firing at low rates (see Roxin et al., 2011 for a theoretical discussion and Osada, Nishihara, and F. Kimura, 1991 for an example) relative to small bin widths (e.g. 10ms). For example, binning at high temporal resolution would result in a large proportion of empty bins. A correlation coefficient computed from these vectors would treat the common bin pair (0,0) as equivalent to the much more meaningful yet infrequent pair (1,1).

One solution may be to simply increase the bin width. However, at a resolution of 100ms, for instance, interspike intervals (ISIs) of 50ms would often be deemed synchronous. Very little information on synaptic architecture - the presumed basis for Hebbian learning (Cooper, 2005) - could be inferred. As ISI values grow, the true size of the inferred assembly grows in tandem with the number of potential synapses involved. Thus, a method with such resolution could only characterize network dynamics involving a very large number of cells over long timescales and provide limited inferences on their connectivity. Thus, binning and correlation analysis is not ideally suited to computation of pairwise synchrony.

Therefore, the need arises to measure pairwise synchrony with greater precision. There is, however, a lower limit to measurable synchrony imposed by neuronal jitter. Conduction rates, neural morphological properties, and noise from nearby cells may all contribute to jitter (Kuriscak et al., 2012; Kilinc and Demir, 2018). It is thus fundamental to aim to detect not pure synchrony (i.e. on a 1ms scale), but to design synchrony detection around what is deemed a ‘permissible’ jitter (Pazienti, Diesmann, and Grün, 2007; Pipa, Wheeler, et al., 2008).

The challenge of defining *δ* is thus one of accurately reflecting the information content of sparse spike trains. One approach, described by Berger, is to simply count the number of coincident spikes *t*_*i,j*_ - allowing for some jitter *ψ* - and subtract from this total the number of expected coincidences 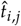 (Berger et al., 2010). The resulting quantity is termed a conditional spike frequency, or CSF. Clearly, such a measure of *δ* is sensitive to the rates of the two neurons. If this were used directly as a basis of computing *ε*, resulting assemblies would be heavily biased towards neurons with high rates. To improve the metric, we choose to compute the gain in firing rate that *i* experiences in the presence of a spike by *j* relative to its mean rate across the recording session:

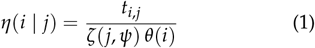

where *ζ*(*j, ψ*) and *θ*(*i*) are the total time during which spikes of *j* occur with jitter *ψ* and the mean rate of *i* across the session, respectively. Noting that the ratio between *t*_*i,j*_ and *ζ*(*j, ψ*) is the firing rate of *i* at times determined by the presence of a spike by *j*, the value of *η*(*i* | *j*) is the ratio between the conditional rate of *i* and its mean rate. Henceforward, we adopt the notation *η*_*i,j*_ for simplicity.

Importantly, the value of *η*_*i,j*_ is purely determined by *t*_*i,j*_ when total spike counts are kept constant. By taking the residual between *η*_*i,j*_ and the mean across the entire population, we now have:

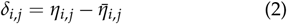

The distribution of *δ* is centered around zero. When *δ*_*i,j*_ < 0, *i* and *j* are less coactive than would be expected from the sampled population. Conversely, when *δ*_*i,j*_ > 0, *i* and *j* are more coactive than expected.

### Significance Testing of *δ*

Testing *δ* for significance is largely a judgement on *η*_*i,j*_ if the population mean can be assumed to be representative. In turn, any significance test on *η*_*i,j*_ is simply a test on the number of coincidences *t*_*i,j*_ if overall spike counts are preserved. We therefore argue that p-values derived for *t*_*i,j*_ are directly transferable to *δ*_*i,j*_.

To this end, many approaches have been used to test the significance of a pairwise coincidence detection metric (Berger et al., 2010). Here, we choose to shift (dither) spikes in both spike trains independently of all others, creating a null distribution of coincidence rates that preserves the rate profile and ISI distribution to a precision determined by the maximum dither width *ω* (Louis, 2010). The dither for each spike is an independent discrete random variable chosen from a uniform distribution 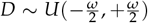. As is often the case, though, the power of surrogate data generation is limited by the number of surrogates, hence computational efficiency.

To overcome this challenge, we derive the analytical distribution of *t*_*i,j*_ given *ψ* following the dithering operation with parameter *ω*.

When evaluating *t*_*i,j*_ for a pair of spike trains, we quantify the number of ISI values *ν* such that |*ν* | ≤ *ψ*. To find *t*_*i,j*_ after dithering, we must understand the effect that dithering both spike trains has on the distribution of *ν*. For convenience, we will refer to the value of *ν* for the *k*th spike in a train after dithering as 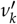. Note that dithering can either preserve, destroy, or generate coincidences: coincidences (|*ν*_*k*_| ≤ *ψ*) may remain or be lost, while noncoincidences (| *ν*_*k*_ | > *ψ*) may remain or be converted into coincidences. Clearly, the probability *ρ* of each of these events depends on *ν, ω*, and *ψ*. As *ω* and *ψ* are constants, only *ν* varies within a dataset. If *ρ*(*ν*) is known, the expected value of *t*_*i,j*_ following dithering may be calculated as the sum of *ρ*(*ν*_*k*_), with the frequency of each value of *ν* essentially representing a set of weights. A similar process is described by Pazienti, Maldonado, et al., 2008.

Crucially, the expected value of *t*_*i,j*_ does not itself represent a test of significance. Rather, the distribution of *t*_*i,j*_ following dithering is required. This distribution *B*(*t*_*i,j*_) is obtained by convolution of the distributions *B*_*ν*_(*t*_*i,j*_) corresponding to each value of *ν*. Each of these is a binomial distribution where the probability of success is defined as the likelihood that 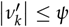 and the number of trials equals the number of spikes with *ν* = *ν*_*k*_. By considering each value of *ν* individually, regardless of whether it repeats, we derive *B*(*t*_*i,j*_) as a Poisson binomial distribution (PBD) where the probability of ‘success’ (i.e. coincidence) is not necessarily constant (Hong, 2013) between trials (i.e. values of *ν*).

We therefore begin by calculating 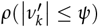 as a function of *ν, ω*, and *ψ*. Here, the probability mass function from which *ρ* is derived is that of the dithering operation defined by *ω*, given that *ν* and *ψ* are known and the only source of variability is the combined effect of independently dithering two spikes according to 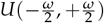. This combined effect is the sum of two uniform random distributions, namely a variant of the triangular Irwin-Hall distribution (Philip, 1927) with *n* = 2. As such, we obtain:

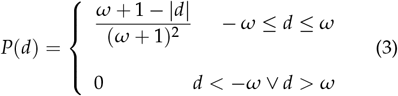

where *d* = *ν*′− *ν* represents the combined effect of dithering two spikes on *ν* (Pazienti, Diesmann, and Grün, 2007; Pazienti, Maldonado, et al., 2008). This distribution *P*(*d*) describes the probability of net shift *d* when two spikes are independently dithered. We then estimate the likelihood of a coincidence being retained or injected as follows:

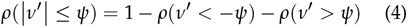

Next, we expand the second and third terms of the above by recognizing that *ρ*(*ν*′ < −*ψ*) = *ρ*(*d* < −*ψ*− *ν*) and *ρ*(*ν*′ > *ψ*) = *ρ*(*d* > *ψ* − *ν*). With Eq.3, we obtain:

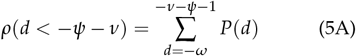

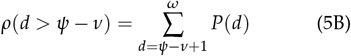

Eq.5A and 5B represent the probability of noncoincidence through violation of the lower and upper bounds −*ψ* and *ψ*, respectively. With this, we state the post-dithering PBD of *t*_*i,j*_:

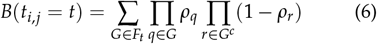

where *ρ*_*k*_ denotes 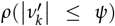, *F*_*t*_ is the set of all subsets of *t* integers that can be selected from {1, 2, 3, …, *n*}, *n* is the number of values of *ν*_*k*_ in the dataset, and *G*^*c*^ is the complement of *G*. Eq.6 is the analytical solution that describes the probability of obtaining *t* coincidences after independent uniform dithering of both spike trains with total dither width *ω* and allowed jitter *ψ*. Other than assuming that each *ν*_*k*_ is affected by dithering independently with respect to all other values of *ν* - which is only an oversimplification if a single spike is part of multiple coincidences prior to or after dithering - this derivation is unbiased towards both firing rate profiles and population dynamics without sacrificing temporal precision. This distribution is key in that it allows for the analytical calculation of p-values from pairs of spike trains at even the smallest bin sizes.

Due to combinatorial explosion in enumerating *F*_*t*_, though, it is not possible to directly compute the cumulative distribution function (cdf). However, algorithms capable of managing this complexity with reasonable run-times and memory requirements have been developed. Henceforward, we adopt a version of the fast Fourier transform algorithm discussed by Hong and implemented in Python by Straka (Hong, 2013; Mika, 2017).

### Defining a Neuronal Assembly

We define an assembly *E* as any group of *n* cells in a population of size *N* ≫ *n* that satisfies three key criteria:

A. *Coactivity*: Its measure of collective coactivity - an assembly coactivity score *ε*_*E*_ - is significantly greater than that of equivalently sized assemblies when selected at random from the recorded population.
B. *Connectivity*: When viewed as an undirected graph with cells as nodes and significant pairwise coactivity values as edges, the graph of *E* is connected.
C. *Irreducibility*: ∄ *n* ∈ *E* | *ε*_*E*−*n*_ > *ε*_*E*_ where *E* − *n* satisfies **A** and **B**, and ∄ *m* ∉ *E* | *ε*_*E*∪*m*_ > *ε*_*E*_ where **A** and **B** are again satisfied by *E* ∪ *m*.

By **A**, we require assemblies to be rare by definition – that is, their measure of coactivity *ε* given *n* must be statistically anomalous when compared to randomly generated assemblies of the same size. Criterion **B** ensures that high coactivity is not simply driven by isolated strong pairwise links, but rather reflects a connected network. Formally, two disjoint subsets *E*_1_ and *E*_2_ may form a valid assembly only if at least one cross-pair exceeds the significance threshold *q*. Put otherwise, given *E*_1_ and *E*_2_ such that *E*_1_ ∩ *E*_2_ = ∅, then *E*_3_ = *E*_1_ ∪ *E*_2_ may only be considered an assembly if:

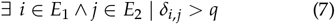

where *q* is the significance threshold for *δ* evaluated between cells *i* and *j*. Essentially, *E* cannot be the union of two smaller assemblies that are not themselves connected by a significant coactivity score between at least one pair of cells. Finally, by **C**, we require that in no case can a cell *m* be added nor a cell *n* removed from *E* such that the resulting assemblies *E* − *n* and *E* ∪ *m* still satisfy **A** and **B** while also exhibiting a greater value of *ε* than *ε*_*E*_.

Intuitively, we expect that as *n* → *N*, an assembly is more likely to satisfy **B** due to the increased likelihood of at least one pair of cells *i, j* satisfying (1). Conversely, **A** is more likely satisfied by assemblies of highly connected cells with *n* ≪ *N*. Henceforward, we refer to a set that satisfies **A** as a putative assembly, a set that satisfies **A** and **B** as a candidate assembly, and a set that satisfies **A, B**, and **C** as a detectable assembly. We use this tripartite definition to draw attention to several key assembly properties of interest, namely:

1. That assemblies are not biologically nor statistically isolated networks;
2. That assemblies should be detectable as foci of pairwise coactivity that exceed what is expected from random sampling of the population;
3. That for an assembly to be detected from pairwise measurements, all member cells should exhibit a degree of connectivity with all other cells, be that direct or indirect;
4. And finally, that assemblies are not necessarily mutually distinct, and there is no reason *a priori* that their member neurons cannot form a limited number of significant connections with non-member cells;

### Assembly Coactivity Scores (*ε*)

The matrix Δ of all pairwise scores *δ*_*i,j*_ is now associated with a corresponding matrix of p-values. When deriving an expression for *ε* from Δ, values that are not significant should be excluded from subsequent analysis. To achieve this, we sparsify Δ by replacing all non-significant values with 0. All *δ*_*i,j*_ for which *t*_*i,j*_ does not show sufficient excess synchrony are thus replaced by a value indicating no deviation from the mean of Δ. To account for multiple comparisons, we choose to apply a Dunn-Šidák correction (Šidák, 1967) factor to *α* = 0.05. However, we choose not to use the number of unique cell pairs as the number of comparisons, as the comparisons themselves are not entirely independent (if *δ*_*i,j*_ and *δ*_*j,k*_ are both known, there is at least some minimal information to infer *δ*_*i,k*_, for example). Rather, we use the number of neurons - fully independent under the null hypothesis - as the correction factor. Thus, we may be confident that all expressions of *ε* are based on pairwise coactivity scores that accurately reflect the measured network. Assuming that most cell pairs do not exhibit significant correlation, the matrix Δ′ obtained by sparsifying Δ will contain relatively sparse non-zero entries. Any method used for clustering cells in Δ′ into assemblies must take this into account.

Δ′ is a subset of all possible positive pairwise connections between cells, and as such may be interpreted as a graph where each pair of cells *i, j* is connected by an edge if and only if *δ*_*i,j*_ > 0. Here, we must pause to recognize that this representation of precise network dynamics is already amenable to a plethora of existing network clustering algorithms (Mölter, Avitan, and Goodhill, 2018; Tavoni, Cocco, and Monasson, 2016; Singh and Garg, 2022; Liu and Barahona, 2020), especially those that allow for detection of overlapping cliques. While we do not intend to explore these options in detail here, we do not in any way exclude the possibility that there already exist methods capable of identifying neural assemblies as defined by **A, B**, and **C** from Δ′. Rather, we present a clustering algorithm premised on the verification of these three criteria via the determination of an assembly coactivity score *ε*, which is the topic of the remaining portion of this section.

First, observe that a reasonable expression of *ε*_*E*_ may take the shape of a linear combination of *δ* ∈ *I* and *δ* ∈ *O*, where *I* is the set of *δ*’s for all unique cell pairs in an assembly *E* and *O* is the set of *δ*’s for all unique cell pairs with only one member in *E*:

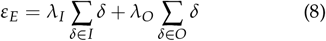

where *λ*_*I*_ and *λ*_*O*_ are weights that apply equally to all elements of *I* and *O*, respectively. Here, *λ*_*I*_ and *λ*_*O*_ determine how much weight is given to internal vs. external coactivity. For assemblies to stand out as foci of coactivity, *λ*_*O*_ must be negative, penalizing external connectivity. In the simple case where we take *λ*_*I*_ = 1 and *λ*_*O*_ = −1, *ε* is reduced to the difference between the sum of all internal pairwise coactivities ∑ *I* and that of neighboring cells ∑ *O*.

Without appropriate normalization, however, larger assemblies are favored simply because they accumulate more edges. For example, if a candidate neuron *m* is equally coactive with two assemblies *E* and *F*, but *F* is slightly larger, then *m* will appear more strongly linked to *F* — biasing the detection process. Thus, *ε* must be scaled by assembly size. We may achieve this by normalizing the first and second terms in Eq.8 by a function of the size of *E* and *E*^*c*^, respectively.

An intuitive normalization procedure would to set *λ*_*I*_ and *λ*_*O*_ to the inverse of the size of *I* and *O*, respectively. Here, *ε* is now the difference between the mean of *I* and that of *O*. Again, though, we encounter a problem when dealing with sparse networks. Should a cell *m* be a plausible addition to *E*, the assembly *F* = *E* ∪ *m* would only satisfy **C** if the addition of *m* could raise *ε*. Even assuming that *m* is as uncorrelated with the rest of the population as every cell in *E*, **C** would not be met if *m* were to have a number of *δ* values shared with other members of *E* that is only marginally below the mean (assuming all connections are approximately equal). In the most absurd scenario, a cell connected to 8 out of 10 cells in *E* would not be considered part of the same assembly if each of those 10 cells were connected with the other 9. Thus, when searching for an appropriate value of *λ*, removing the size bias from *ε* may not be entirely desirable.

As we have considered and rejected two extreme values for *λ*, we argue that defining *λ*_*I*_ and *λ*_*O*_ as the inverse of the number of neurons in *E* and *E*^*c*^ respectively is sufficiently permissive to both regular and overlapping assemblies. We thus propose:

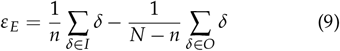

where *N* is the size of the population and *n* is the size of *E*. Henceforward, we adopt Eq.9 as our definition of *ε*_*E*_. This formulation ensures that assemblies are defined as groups whose mean internal coactivity exceeds their mean external coactivity, while remaining robust to both sparse connectivity and overlapping membership.

### Assembly Detection Algorithm

Here, we describe an algorithm that iteratively scans the population for assemblies of a given size *n*, then filters these assemblies using criteria **A** and **B**. The filtered group of candidate assemblies is then used to generate a possible set of derivative assemblies of size *n* + 1 via the addition of a single neuron. More precisely, the steps of the algorithm are as follows:

**Step 1** *Initializing triplets* We compute *ε* for all possible assemblies of size three. We filter these assemblies by verifying criteria **A** and **B**.

**Step 2** *Assembly derivation* For each putative *E*, we generate derivative assemblies by adding each neuron from *E*^*c*^ in turn. All combinations are produced.

**Step 3** *Permutation filtering* To avoid redundant testing, assemblies that differ only by the ordering of their members are collapsed into a single candidate.

**Step 4** *Verifying coactivity* To verify criterion **A** without knowing the full distribution of *ε*, we sample random assemblies of the same size to derive an estimate for the mean and standard deviation of *ε*. From these, we obtain a conservative lower bound for significant *ε*. Assemblies that meet or exceed this bound are deemed putative assemblies.

**Step 5** *Verifying connectivity* Putative assemblies are filtered by only taking those that satisfy criterion **B**. This produces a set of candidate assemblies. **Step 6** *Iteration* Steps 2-5 are repeated until either a maximum assembly size is reached, or the set of candidate assemblies is empty.

**Step 7** *Verifying irreducibility* For each thus identified assembly, criterion **C** is verified. Assemblies that pass this final filter have been detected by the algorithm.

**Step 8** *Superset filtering* When assemblies overlap, it is likely that their union is also detected as a third assembly. These supersets are removed.

This algorithm is efficient in that, for each assembly size, only assemblies that are more likely to pass **A** and **B** than an assembly selected at random are tested. Efficiency arises because only assemblies with significant precursors are tested at larger sizes: if *ε*_*E*_ is not significant, then *ε*_*E* ∪ *m*_ is unlikely to be significant for any *m*.

As we do not exhaustively enumerate all assemblies of size *n*, we cannot directly compute the null distribution of *ε*. Instead, we estimate its mean *µ* and standard deviation *s* from a random sample of assemblies (Gurland and Tripathi, 1971). Applying the Chebyshev inequality (Chernick, 2011), we set a conservative threshold: the probability of a random assembly exceeding *k* standard deviations above the mean is at most 1/*k*^2^. This method provides a nonparametric, conservative significance threshold.

### Evaluating Clustering Performance

We employ a modified version of the Rand index (RI) and adjusted Rand index (ARI) to quantify the agreement between programmed assemblies and detected assemblies (Hubert and Arabie, 1985). RI computes a pairwise measure of agreement as the fraction of true positives (TP) and true negatives (TN) across all cell pairs:

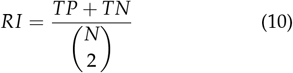

where TP is the number of pairs *i, j* that appear in the same cluster in both the set of ground truth (GT) clusters and that of detected clusters, and TN is the number of pairs *i, j* that appear in different clusters in both the set of GT clusters and that of detected clusters. Importantly, the set of cells that are not part of any assembly constitute a cluster in both GT and detected datasets.

RI is bound by 0 and 1, with extremes representing complete disagreement and complete agreement between GT and detected assemblies, respectively. ARI, instead, accounts for the expected number of TP and TN as follows:

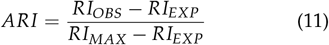

where *obs, exp*, and *max* denote the observed, expected, and maximum possible value of *RI*. Here, we set *RI*_*MAX*_ = 1, as we believe that perfect clustering should be achievable. We approximate *RI*_*EXP*_ ≈ 0.5, which is closely matched by simulation results. This implies that cell pairs sampled from random clusters generated with the same size and number of overlapping elements as those actually detected have a 50% chance of being arranged as in GT. The bounds of ARI are therefore −1 ≤ *ARI* ≤ 1, where the extremes represent complete disagreement and complete agreement, respectively. An ARI value of 0 indicates performance at chance level (i.e. the algorithm performs as it would if it only had access to pairwise measurements but no information on their clustering properties).

RI and ARI are conventionally used to evaluate the performance of algorithms that do not permit overlapping clusters (as these are typically not present in GT). Therefore, we must clarify that our definition of TP and TN includes all clusters in which the two cells in question appear. Put otherwise, two neurons are considered to agree if they co-occur in at least one cluster, rather than in the exact same cluster. In practice, this modification is permissive to overlapping cells but does not significantly impact overall performance if the number of cells belonging to more than one assembly is ≪ *N*.

### Assembly Pattern Vectors

A key aim of assembly analysis is the discovery of a pattern vector for each detected assembly (Lopes-dos-Santos, Ribeiro, and Tort, 2013). The assembly pattern vector *W* is a set of weights that allows the activity of the assembly to be tracked over time as follows:

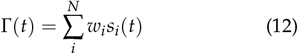

where *w*_*i*_ and *s*_*i*_(*t*) are the assembly pattern weight associated with neuron *i* and the number of spikes of neuron *i* at time *t*, respectively. To determine *W*_*E*_, we evaluate the contribution of each neuron to *E*:

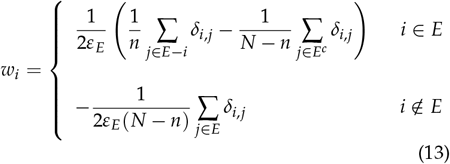

Eq. 13 assumes equal contributions of two neurons to their pairwise coactivity score. When a neuron is a member of the detected assembly, its weight is therefore the difference between what coactivity it shares with the rest of *E* and what it shares with *E*^*c*^. Conversely, every neuron that is not a member of the detected assembly only contributes to values of *δ* in *O*, not *I*. Further, observe that ∑ *w*_*i*_ = 1 (see Appendix A), and that each *w*_*i*_ is normalized by *ε*. This ensures that the range of values for Γ(*t*) is similar for all assemblies.

### Simulation Methodology

All simulations, unless otherwise noted (e.g. Fig. 1), comprise 50 neurons modeled as Poisson processes for a 30 minute recording. The non-zero rate of each neuron is itself drawn from a Poisson distribution (*λ* = 3). Bins are 1ms in width, and can accommodate no more than one spike per neuron.

**Figure 1:**
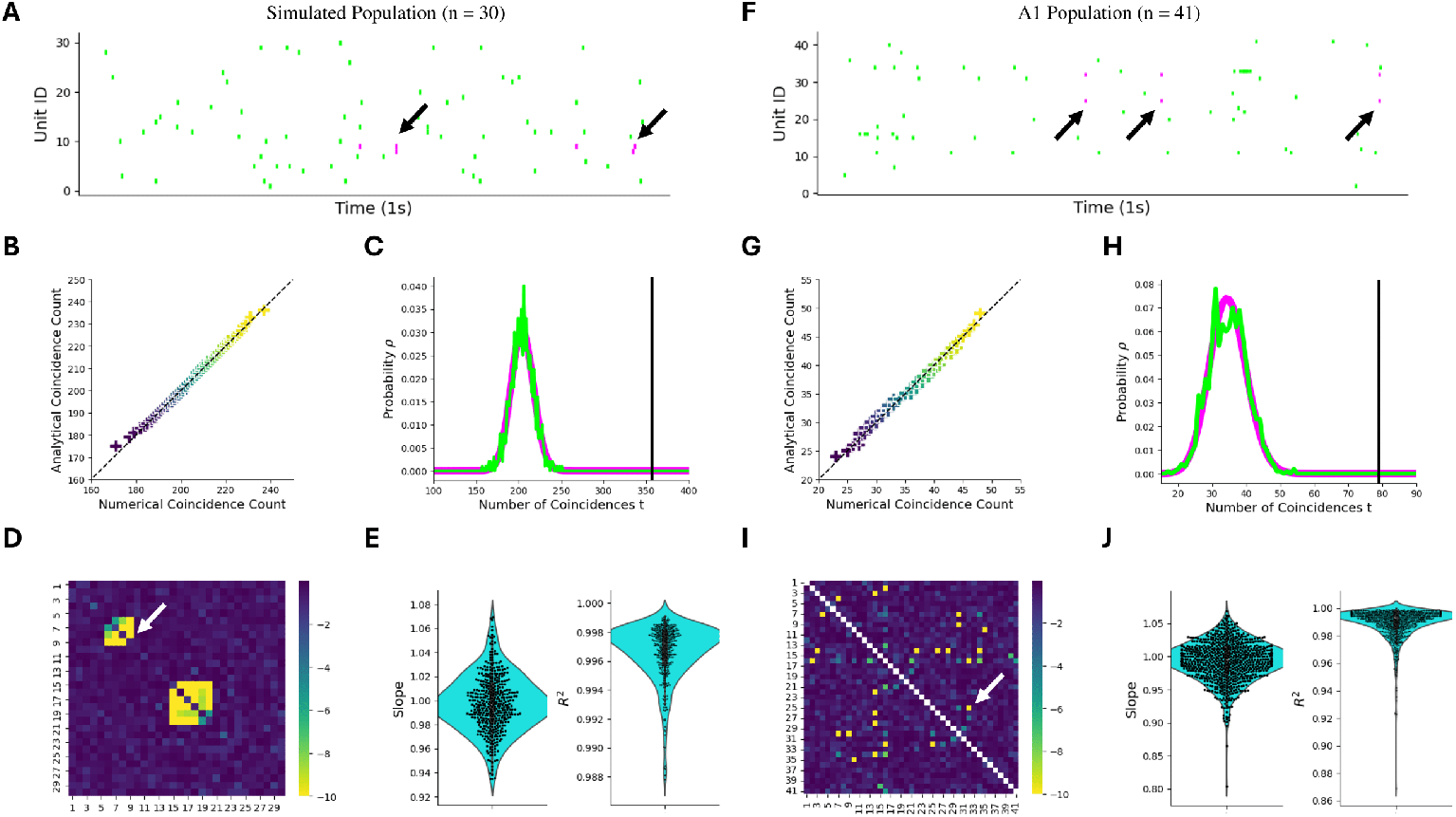
Validation of pairwise synchrony analysis in simulation (left panels) and an example A1 recording (right panels). A-F) Raster plots (1s segments) with a pair of synchronous neurons highlighted in purple. Arrows point to coincident firing. B-G) QQ plots of the distributions shown on the right, comparing the numerical and analytical coincidence counts (<=5ms intervals). Numerical coincidence counts are produced through generation of 1000 surrogate datasets by dithering (25ms). C-H) Coincidence counts probability-mass functions (PMFs) of numerical (green) and analytical (purple) methods, with the observed number of coincidences indicated by the black vertical line. In both cases, this pair of neurons is clearly synchronous. D-I) Matrix of analytical p-values: each entry i, j is the negative base-10 logarithm of the corresponding p-value of δ_i,j_. White arrows indicate the pair shown in the panels above. For the simulated population, clustering of hidden processes E_1_ and E_2_ is evident. E-J) Linear regression results from analysis of QQ plots. The distribution of best-fit slopes is shown on the left, with that of R^2^ values shown on the right.

Assemblies are modeled as hidden processes. Here, a ‘hidden’ mother neuron is modeled as a Poisson process with a 2Hz rate. All neurons selected to be part of the corresponding assembly receive copies of the spikes from the hidden process with a certain probability *ϕ*(*λ*), where *λ* is the rate of the receiving neuron. To scale *ϕ* appropriately, we use:

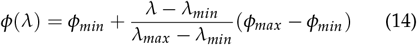

where *ϕ*_*min*_, *ϕ*_*max*_, *λ*_*min*_, and *λ*_*max*_ are the minimum and maximum copy probabilities and rates across the simulated population, respectively. With Eq.14, the copy probability is linearly scaled to match the natural rate of the receiving neuron. For consistency, we always take *λ*_*min*_ = 1*Hz* and *λ*_*max*_ = 10*Hz*. This imposes an upper bound on *ϕ*_*min*_ of 0.1. Due to the rate of the hidden process being capped at 2Hz, we therefore ensure that the activity of neurons participating in a single assembly depends on the hidden process for at most 20% of all spikes. Therefore, an average 3Hz process will generate 5400 spikes across 30 simulated minutes, and, if participating in an assembly, about 1080 more.

### Decoding Methods

Receiver operating characteristic (ROC) analysis was used to measure rate-based decoding power. Formally, decoding power was quantified as:

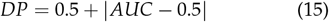

where AUC is the area under the ROC curve. Briefly, ROC curves plot the true-positive rate against the false-positive rate obtained from a binary classifier. The greater the AUC, the greater the ability of this classifier to differentiate between two groups. For decoding analysis, these groups were defined as stimuli that induced the greatest response in the decoded cells (calculated as the mean across trials). Response itself was measured in one of two ways, yielding two forms of rate-based decoding:

1. The total number of spikes produced by the decoded population during stimulus presentation (pure rate code);
2. The total number of coincident spike pairs produced by the decoded population during stimulus presentation (coincidence or synchrony-based decoding);

The latter code emphasizes the importance of synchrony without explicitly requiring consistent temporal organization across trials.

Temporal codes were also analyzed, this time by binning assembly spike trains and computing their correlation matrix across all repetition trials. The decoded stimulus was defined as that to which the response was most correlated on average.

## Results

Data was obtained either by simulation (see Simulation Methodology) or from ferret auditory cortex during passive listening (see Pennington and Stephen V. David, 2022). Simulations were used in conjunction with ferret data to validate assembly detection methods. These methods were then applied to recorded populations in the ferret to study and decode network activity. As the original dataset included both unique and repeated stimuli, decoding was attempted on repetition trials (20 per each of 18 stimuli per session). Assemblies, though, were computed irrespective of stimulus repetition count.

### Detection of Pairwise Synchrony

To validate the analytical method for detecting pair-wise synchrony, all pairs of <10Hz units (n = 41 units, n = 820 unique pairs) from a single A1 recording session were evaluated both analytically and numerically (Fig. 1, right). The same procedure was applied to a simulated recording (n = 30 units; Fig. 1, left). Maximum dither was 25ms, and maximum ISI for synchronous spikes was 5ms. 1000 surrogate datasets were produced for each validation experiment.

Then, for each pair of units, percentiles of coincidence counts were computed from both the numerical probability mass function (PMF) and analytical PMF. Representative examples of both PMFs are shown in Fig. 1C and 1H (purple denotes observed number of coincidences), with corresponding percentiles plotted against each other in Fig. 1B and 1G, respectively. From plots like that in Fig. 1B, we extract the slope and R-value through linear regression. The full distributions of both metrics are plotted in Fig. 1E and 1J. By median, analytical results explained 99.3% of variance in numerical results, indicating near-perfect modeling in the A1 dataset. This figure closely matches that achieved in simulations (99.7%). Outliers were examined visually for potential biases. All such cases appear to have been due to the discrete nature of coincidence counts, which is reflected in the percentiles of numerical PMFs. Thus, generating a greater number of surrogates and computing more quantiles would be expected to resolve remaining inconsistencies.

### Assembly Identification

We next sought to determine the efficacy of the assembly detection algorithm (see Methods). As ground truth is not available in ferret datasets, we performed this analysis in simulations only. To this end, we conducted two experiments (data in Fig. 2A-H). First, we performed 20 simulations (*ϕ*_*min*_ = 0.1) of three partially overlapping assemblies: *E*_1_ = {6, 7, 8, 9}, *E*_2_ = {9, 19, 20, 21, 22, 23}, and *E*_3_ = {23, 32, 33, 34, 35, 36, 37, 38, 39}. In just one sim-ulation, cell 9 was not identified as a member of *E*_2_.

**Figure 2:**
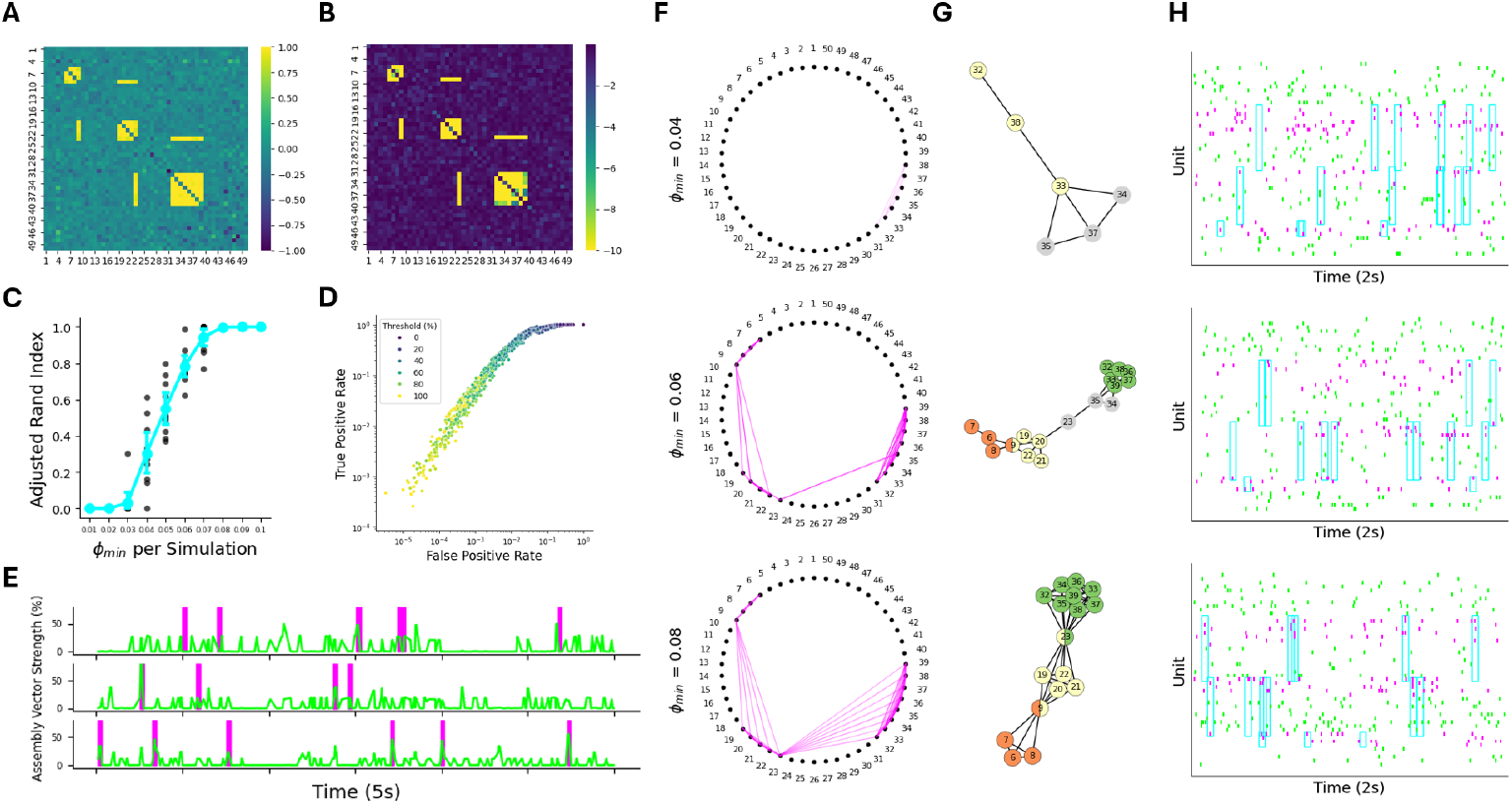
Three-assembly simulations with detection and activation analysis. A) Example Δ matrix of pairwise coactivity scores. B) The corresponding p-value matrix (negative logarithm transform) associated with A. C) ARI with varying minimum copy probability, showing optimal assembly detection with ϕ_min_ > 0.07. 10 simulations were conducted per value of ϕ. D) Log-log ROC curve from 20 simulations with ϕ_min_ = 0.1. Color warmth is mapped to % assembly strength threshold, used to infer activity of the hidden process. E) An example 5-second time series of inferred assembly activity (E_1_ top, E_2_ middle, E_3_ bottom) with ground truth activity highlighted as purple 20ms bins. ROC analysis was performed by applying thresholds to assembly pattern vectors such as these. F) Example polar plots for varying ϕ_min_, with connections between nodes representing synchrony between cell pairs. G) Example graph diagram of assembly cells, shown as nodes with significant pairwise synchrony scores δ as edges. Colors indicate the detected parent assembly. Gray nodes were not identified as part of any assembly. H) 2-second raster plots of simulations used to test the limits of assembly detection through reduction of ϕ_min_ (summary in C). Spikes from assembly member neurons are shown in purple, with cyan boxes showing programmed activation of the hidden process. Note that with higher copy probabilities, these boxes contain a greater proportion of assembly spikes.

Otherwise, all assemblies were correctly identified. Examples of Δ and the corresponding p-value matrix are shown in Fig. 2A and Fig. 2B, respectively. In Fig. 2E, we show example assembly activation traces calculated from their corresponding pattern vectors. Purple-shaded regions identify programmed hidden process activity. To quantify the accuracy of assembly activation traces in representing assembly activity, we performed ROC analysis to decode assembly activity (Fig. 2D). While performance generally increased with assembly size, optimal thresholds could detect approximately 80-90% of assembly activity with a false-positive rate of circa 20%.

Second, we searched for the lower bound on hidden process activity for assembly detection. We performed 100 simulations with 10 different values of *ϕ*_*min*_, ranging from 0.01 to 0.1. Results are shown in Fig. 2C and examples in Fig. 2F-H. With increasing copy probability, we observed improved assembly detection as quantified by the adjusted rand index (ARI; see Methods). This improvement was most significant between *ϕ*_*min*_ = 0.04 and *ϕ*_*min*_ = 0.08 (Fig. 2C). In the latter case, over 93% of assemblies were detected. Crucially, the value of *ϕ*_*min*_ can be easily translated to the percentage of spikes originating from the hidden process. A 3Hz cell with *ϕ*_*min*_ = 0.08 corresponding to a 2Hz hidden process will participate in the assembly at a rate of approximately 0.48Hz (due to *ϕ* scaling and the assembly rate). This represents an additional 16% of activity above baseline, or about 13.8% of total activity per cell. The likelihood of two or more assembly cells being active simultaneously is lower still. Therefore, we argue that this assembly detection method is sufficiently sensitive even at low copy probabilities and consequently resistant to large volumes of uncorrelated (noisy) activity.

### Synchrony or Correlation?

Following validation experiments, we performed assembly analysis of ferret datasets to characterize network activity. To ensure that assemblies were not skewed by small population sizes, we performed all subsequent analysis on recording days with a minimum of 20 units. As excess synchrony could in principle arise from trial-by-trial pairwise correlation, we compare both metrics across neuron pairs either included or excluded from identified assemblies in A1 (Fig. 3). The equivalent process in PEG is shown in Fig. S1. Here, Fig. 3A shows an example trial with spikes from assembly member units in purple (cyan box).

**Figure 3:**
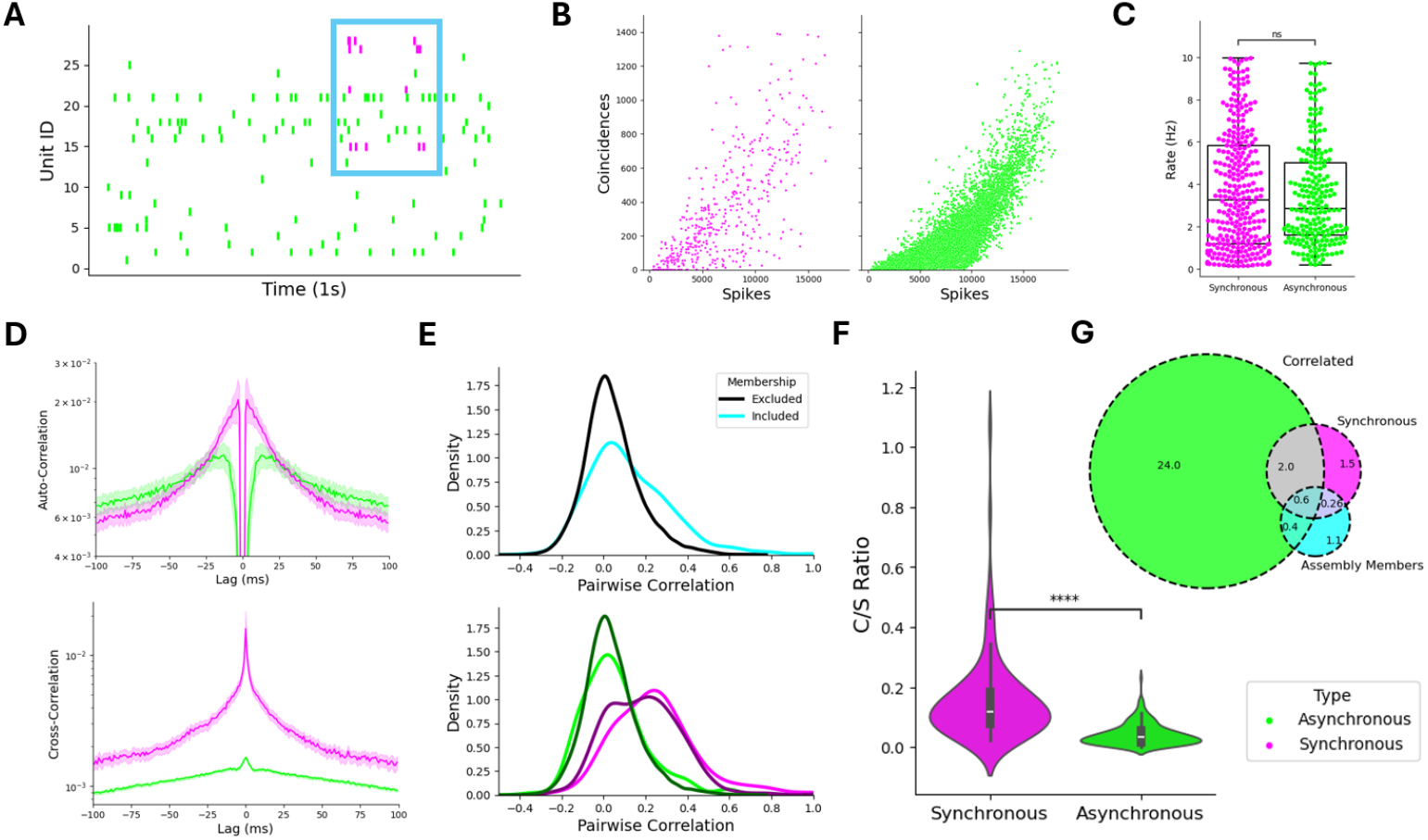
Adirectional pairwise temporal dynamics in synchronous and asynchronous units. A) Example raster plot from A1 recording session, with one four-unit assembly shown in purple. The assembly appears to be synchronously active twice in this window, with all units participating (cyan box). B) Coincidence counts and spike counts for synchronous (purple) and asynchronous (green) pairs. Observe that, on average, synchronous pairs generate more coincident events per spike. C) Distribution of mean firing rates for neurons synchronous with at least one other unit (left) and those that are not synchronous with any other cell (right). D) Auto-correlogram (top) and cross-correlogram (bottom) for synchronous (purple) and asynchronous (green) units. Synchronous units are defined as in C for the auto-correlogram. E) Distribution of trial-by-trial pairwise correlation between pairs included in detected assemblies (cyan) and those that remain excluded (black). The bottom panel further differentiates between synchronous and asynchronous pairs (dark for excluded, light for included). F) Distribution of coincidence counts per spike (C/S) ratio, quantifying the pairwise relationship shown in D. G) Venn-diagram showing pairs of neurons that are correlated (green), synchronous (purple), and included in assemblies (cyan). Numbers indicate the % of all pairs that fall into a given category.

Several important observations stem from this analysis. First, fewer spikes are required to generate coincident events for synchronous pairs (Fig. 3B) compared to asynchronous pairs. Indeed, the ratio between coincidence counts and spikes (C/S Ratio) is significantly greater for synchronous pairs than asynchronous pairs (Fig. 3F; U = 1.634 · 10^4^, p = 7.632 · 10^−27^ two-sided Mann-Whitney test). Second, we observe that there is not a significant difference between the mean firing rates of neurons that do not participate in synchrony compared to those that do (Fig. 3C; U = 3.139 · 10^4^, p = 0.7561 two-sided Mann-Whitney test).

Third, Fig. 3G shows that 27% of all neuron pairs exhibit significant pairwise correlation, while only 4.4% are synchronous (identical significance thresholds employed for both metrics). Synchrony is therefore a property of neural activity that is more sparsely distributed compared to trial-by-trial (i.e. second-by-second) correlation. Fourth, note that the odds of a cell pair belonging to an assembly are about 1 in 42. If the cell pair is synchronous, the odds of belonging to an assembly are 1 in 5. If the cell pair is correlated, its odds are 1 in 27. Thus, odds increase by factor of 8 for synchronous pairs but only by 1.5 for correlated pairs. This suggests that detected assemblies are primarily based on synchrony, not correlation (as intended by Methods). Henceforward, we refer to these as synchrony-driven assemblies.

Fifth, while pairs of neurons included in detected assemblies tend to exhibit greater correlation (Fig. 3E, top), this effect is entirely driven by synchronous pairs (Fig. 3E, bottom). Sixth, both auto-correlograms and cross-correlograms differ between asynchronous and synchronous neurons (Fig. 3D). In the autocorrelogram, units involved in synchrony with at least one other unit demonstrate an increased tendency for burst-firing at short intervals. The difference between cross-correlograms for synchronous vs. asynchronous pairs, however, is far greater: synchronous pairs not only demonstrate consistently greater correlation across intervals, but also produce a much larger central peak. This suggests that spike dithering adequately captures synchrony as evident from cross-correlograms, and that this property may arise from temporal spike clustering. All other cell pairs appear only weakly co-modulated.

### From Synchrony to Sequences

Thus far, we have only considered synchrony without regard for directionality. As temporal firing patterns at short timescales would provide further evidence for the existence of coordinated networks, we searched for robust firing sequences across each sampled population. First, we used a simple binomial test to determine whether unit A tended to fire in the 5ms before or after unit B. Results from a representative population are shown in Fig. 4A, where nodes represent neurons and arrows are drawn from the Pre to Post cell only when the pair was both synchronous and temporally coordinated. Assembly member neurons are shown in purple: their position at the center of this network is indicative of strong temporal correlations with non-assembly neurons. Further, we observed that the proportion of unit pairs that were significantly coordinated was much greater when units were synchronous rather than asynchronous (Fig. 4E; T = 0, p = 4.883 · 10^−4^ two-sided Wilcoxon test). Moreover, an ordered cross-correlogram of only synchronous pairs (Fig. 4F) was clearly indicative of a pre-post relationship. The difference between coordinated (purple) and uncoordinated (green) pairs was small but consistent, with uncoordinated pairs exhibiting weaker cross-correlograms despite optimal ordering. Note that Fig. 4F includes the mirrored cross-correlogram (shadows) to facilitate prepost comparisons.

**Figure 4:**
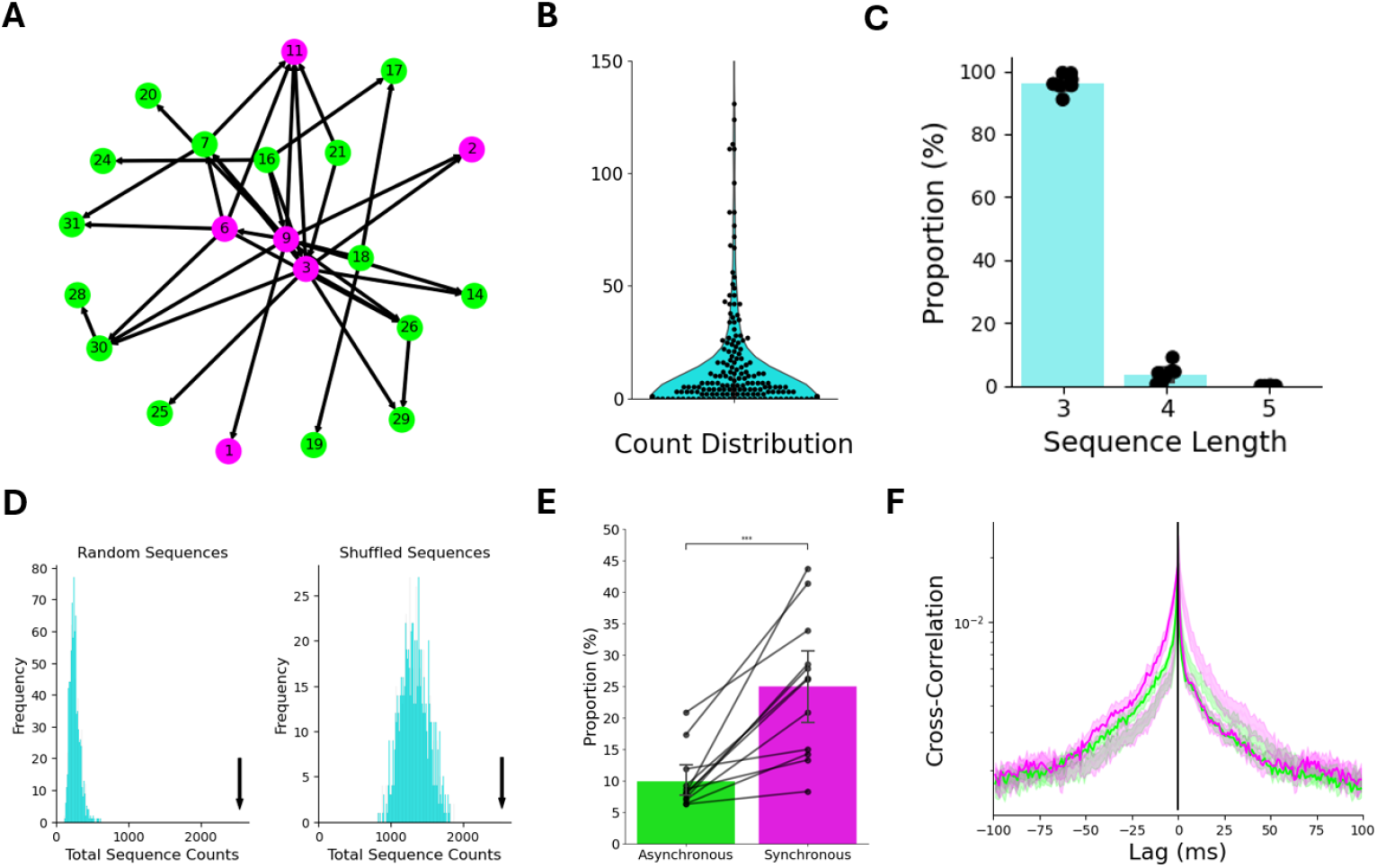
Pairwise temporal dynamics in synchronous and asynchronous units. A) Network representation of an example A1 session, with arrows connecting pairs of units that are both synchronous and temporally coordinated (from pre to post). Assembly members are shown in purple. B) Frequencies of predicted sequences, with predictions made by following ‘paths’ on graphs like those in A. Data from all sessions with at least 5 predictions is shown. C) Proportion of identified predicted sequences by size, per session. The overwhelming majority are short (3 units), with 1-10% consisting of four units and the remainder five. D) Example null distributions of total sequence counts for either random sequences (left) of shuffled predicted sequences (right), generated from the same session as A. In both cases, predicted sequence counts are significantly greater than those of surrogates. E) Proportion of asynchronous or synchronous cell pairs that exhibit significant temporal coordination at 5ms intervals (defined by the pre-post binomial test). Data is shown per session. F) Cross-correlogram for ordered coordinated (purple) and non-coordinated (green) synchronous pairs, for all sessions. The mirror image of each correlogram is plotted as a ‘shadow’ reflected over the black line to facilitate visual comparisons.

We therefore hypothesized that the existence of robust pairwise temporal patterns may form the basis of longer sequences. To test this, we counted all sequences of spikes (from 3-6 cells) with consecutive pairs separated by at most 5ms. We predicted that sequences traced from networks like those shown in Fig. 4A (‘paths’ along the graph) would be signifi-cantly more common than both random sequences and shuffled counterparts (see corresponding examples of null distributions of coincidence counts in Fig. 4D). Indeed, we found that predicted sequences were significantly more common than surrogates (p < 10^−2^ for random surrogates and p < 0.05 for shuffled surrogates in all sessions), showing that pairwise relationships are sufficiently strong to produce longer temporal structures. Predicted sequences occurred with some regularity: the distribution of sequence counts is shown in Fig. 3B. For each recorded population, we also calculated the proportion of expected sequences of a given length (Fig. 4C). The overwhelming majority consisted of just three neurons, with sequences of four being rare (1-10%) and those of five only occurring sporadically. In sum, this analysis shows that synchronous neurons produce stereotypical temporal patterns, both through pairwise order and longer sequences. The PEG dataset resulted in similar observations (Fig. S2) These results support the inference that assemblies are temporally organized, possibly with significant intraconnectivity.

### Assembly Rate Codes

Given strong evidence for assembly-like synchrony and temporal dynamics, we wondered whether and how assemblies may encode the identity of complex stimuli. Thus, we investigated the information content of these assemblies (violet ‘Assembly’ in Fig.5). As a control, 100 surrogate assemblies were randomly generated per identified assembly (green ‘Random’ in Fig. 5). Units in the identified assembly were excluded from this process. The number of units was kept constant.

**Figure 5:**
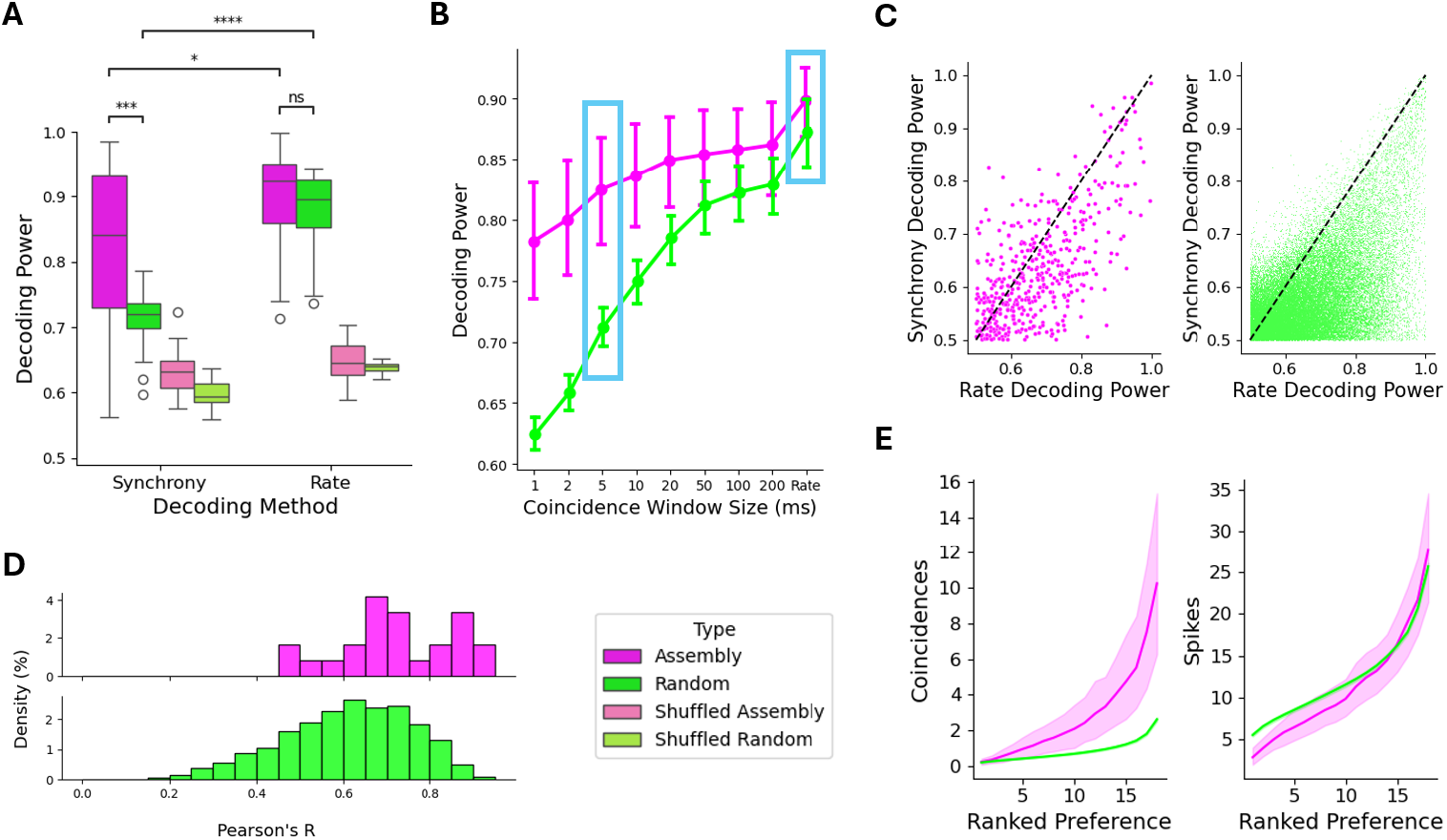
Synchrony-driven assemblies preserve information from rate codes on millisecond timescales in A1. A) Results of ROC decoding analysis, with the area under the curve (AUC) plotted as decoding power for the best decoded stimulus per assembly. Decoding from assembly (n = 26) responses is shown in purple, with surrogates plotted in green. 100 random surrogates were generated per identified assembly. Pale colors show results from trial-shuffling. B) Mean decoding power (AUC) for preferred stimulus with varying maximum interval. Cyan boxes indicate ‘Synchrony’ and ‘Rate’ data shown in panel A. C) Comparison between spike count (rate) and coincidence count (synchrony) decoding for assemblies (left) and random surrogates (right). Dashed lines indicate the first quadrant bisector. Data from all decoded stimuli is shown. D) Distribution of Pearson’s correlation between coincidence count and spike count responses. E) Mean number of coincidences (left) or spikes (right) per stimulus, ordered by response strength per assembly.

Then, the information content of each assembly was assessed during repetition trials by either computing a synchrony score (# of pairwise coincidences from cells in the assembly) or a rate score (total # of spikes produced by cells in the assembly). For each assembly-stimulus pair, ROC analysis was used to evaluate the ability of a binary classifier to decode the presence of the stimulus from either the synchrony score or rate score vectors. The area under the curve (AUC) from each such pair was thus used as a measure of decoding power (see Methods).

As shown in Fig. 5A, the decoding power of the binary classifier varied according to both the metric employed for decoding (5ms-tolerant coincidence count for synchrony and spike count for rate) and the group type (Assembly or Random). Overall, rate was a better predictor of preferred stimulus identity than synchrony (U = 2.140 · 10^2^, p = 2.381 · 10^−2^ for assemblies; U = 1.800 · 10^1^, p = 4.998 · 10^−9^ for surrogates; two-sided Mann-Whitney tests). However, the difference between the predictive power of synchrony-driven assemblies compared to random assemblies was greatest when coincidence count (synchrony) was chosen as the predictor (U = 5.370 · 10^2^, p = 2.804 · 10^−4^ for synchrony-based decoding; U = 4.290 · 10^2^, p = 9.767 · 10^−2^ for rate-based decoding; two-sided Mann-Whitney tests). We also observed that decoding power from shuffled trials (pastel shades) was minimal (<0.7). Taken together, these observations suggest that synchrony-driven assemblies have the unique ability to preserve rate-based information on short timescales.

Likewise, we found that the correlation between assembly responses measured by rate or by synchrony (Fig. 5D) was greater for synchrony-driven assemblies than random assemblies (U = 1.584 · 10^7^, p = 1.788 · 10^−62^; two-sided Mann-Whitney test). Further, the proportion of assemblies better at decoding any - not just preferred - stimulus identity from synchrony than rate was greatest in synchrony-driven assemblies (roughly 30%) compared to random assemblies (20%; Fig. 5C). Thus, groups of synchronous neurons outperform random assemblies in synchrony-based decoding, both when considering their preferred stimulus and when decoding all stimuli.

Generally, inter-stimulus response variance is positively correlated with information capacity as responses become more distinct, hence more amenable to decoding. To test whether this may be a feature of coincidence counts, we ranked stimuli in order of either elicited cumulative spike count or coincidence count per assembly (Fig. 5E). As expected, synchrony-driven assemblies showed a much wider range of coincidence counts per trial compared to random assemblies. Conversely, spike count profiles were very similar between assembly types, implying that mean firing rates are not a significant confound. This result suggests that the firing rates of informative and uninformative units may appear similar despite the presence of cross-neuronal correlations.

Finally, if coincidences but not spike counts truly characterize informative assemblies, we wondered at what time scale this effect may be lost. To investigate this, we performed ROC-based decoding analysis at multiple time scales (1ms – 200ms) for both synchrony-driven and random assemblies (Fig. 5B). Increasing the coincidence window led to a monotonic increase in decoder performance. However, the magnitude of the additional information conferred by synchrony decreased monotonically. Thus, while any assembly will generally provide more information when interrogated at longer time scales, non-spurious responses by synchrony-driven assemblies are most strongly separable within a few tens of milliseconds. Again, similar results were obtained from PEG (Fig. S3).

### Assembly Temporal Codes

As a coincidence-based rate code proved viable in synchrony-driven assemblies, we hypothesized that synchrony-driven assemblies could respond with a reliable temporal architecture to specific stimuli, possibly forming the basis of a temporal code that would be absent in random assemblies. To test this hypothesis, we binned assembly spike trains (combining member neuron spike trains, see diagram in Fig. 6D) and computed the trial correlation matrix, ordered by stimulus identity (Fig. 6A and Fig. 6B). For each trial, we attempted to decode stimulus identity by choosing that to whose responses it was most strongly correlated. We compared the accuracy of this process to that of randomly generated assemblies (with matched size). We observed that both random and synchrony-driven assemblies always exceeded chance level (5.6%), but synchrony-driven assemblies performed significantly better than random counterparts (Fig 6C; T = 69, p = 5.613 · 10^−3^ two-sided Wilcoxon test).

**Figure 6:**
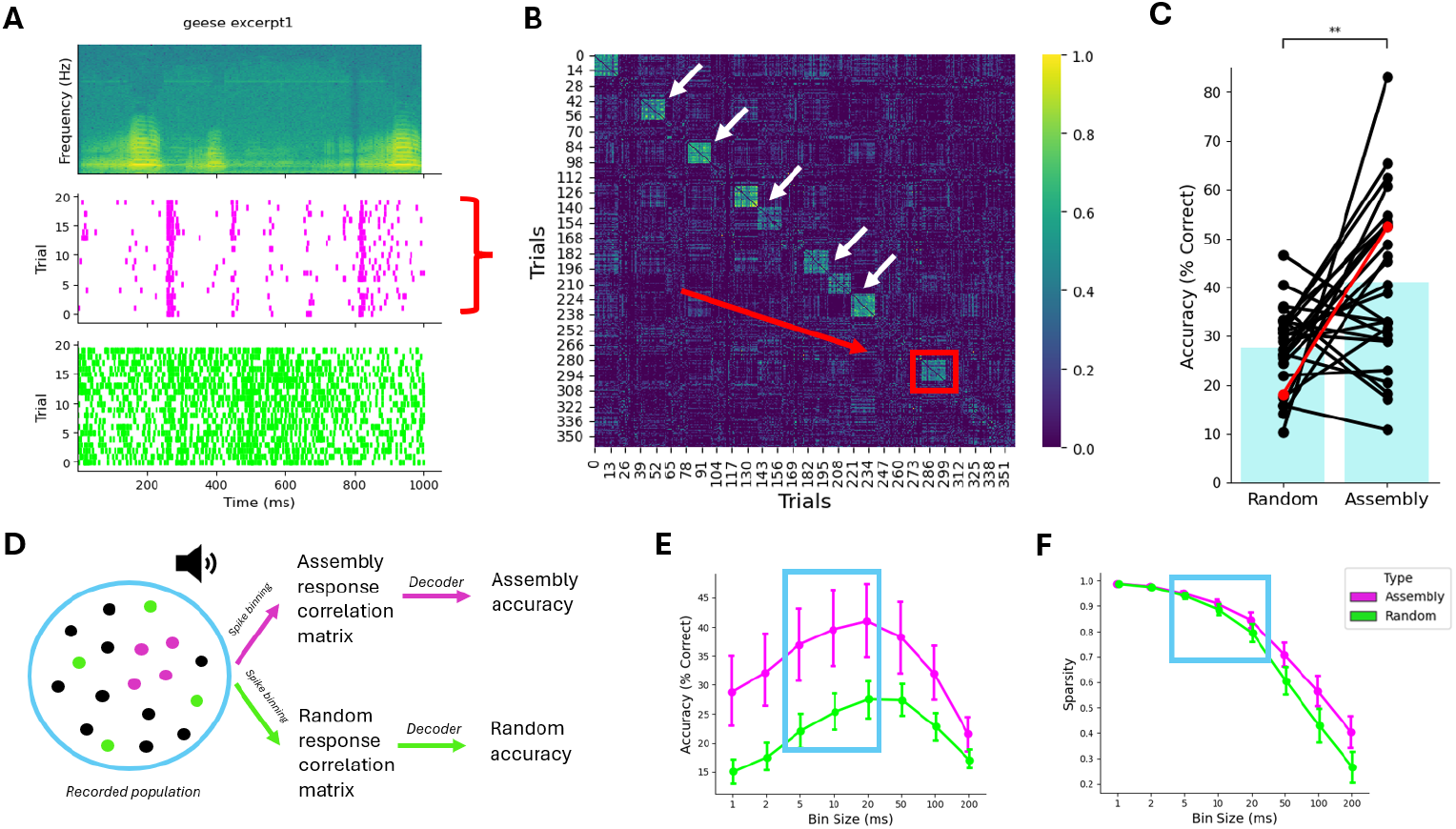
A1 synchrony-driven assemblies employ sparse temporal codes. A) An example stimulus spectrogram (top) with associated trial-by-trial raster plots for one assembly (middle) and all other units (bottom). B) Correlation between assembly response vectors, binned at 20ms, for all repetition trials from one recording session. The example in A is marked in red. Note that trial order is not chronological, and has been rearranged to highlight consistent responses (white arrows). C) Temporal decoding accuracy for assembly-surrogate pairs (n = 26). The example from A is marked in red. D) Diagram of temporal decoding, with assembly and surrogate spike trains being used to generate a correlation matrix that is then decoded to determine accuracy (see Methods). E) Temporal decoding mean accuracy with varying bin width. F) Sparsity - defined as the proportion of empty bins - for assemblies (purple) and surrogates (green) with varying bin width.

As this analysis was performed with 20ms bins, we next tested how decoder performance depended on bin width. Results from varying widths are shown in Fig. 6E, along with the associated spike train sparsity in Fig. 6F. As suggested by rate-based decoding analysis, temporal decoding worked best at bin sizes of 5-20ms. At greater widths, sparsity declined significantly and was always weakest in random assemblies. These results indicate that synchrony-driven assemblies may employ a temporal code with integration constants of 5-20ms, and that sparse activity may improve reliability. We obtained equivalent results in PEG (Fig. S4).

Significant observations were also made on a case-by-case basis: while some assemblies reliably encoded many stimuli (in several cases, most; see an example in Fig. S5), others did not perform as remarkably. Further, different assemblies appeared to encode different aspects of the same stimuli (Fig. S6), implying that a global code could employ parallel synchrony-driven temporal codes to ‘tile’ the response. Ideally, such a prediction would be tested by comparing within-subject assembly responses, but the generally low assembly count prevented this in the ferret dataset.

## Discussion

In these experiments, we have tested the feasibility of conducting pairwise synchrony measurements at a fine temporal scale through an analytical process, employing these metrics as means of assembly discovery, and characterizing the information content of synchrony-driven assemblies in ferret cortical networks exposed to complex natural sounds. Overall, we note that pairwise measurements are very consistent, and that assembly detection, though limited by simulated parameters, can be similarly consistent. Importantly, the ability of synchrony-driven assemblies (SDAs) to better encode the identity of complex sounds through either rate codes or temporal codes - compared to random assemblies - reveals a functional role for synchrony-driven assemblies in cortical sound representation. The similarity between auditory representations in primary and non-primary auditory cortex further characterizes this organization from a systems perspective, and is indicative of shared processing strategies across cortical areas. We discuss these observations in more detail in the following sections.

### Surrogate Data

The effectiveness of dithering as a method for destroying fine temporal structure while preserving firing rate profiles has been thoroughly evaluated in other published work (Pazienti, Diesmann, and Grün, 2007; Louis, 2010; Shmiel et al., 2006). These analyses have shown that dithering can smooth firing rate profiles, but the magnitude of the deformation depends on dither width. We take the view that a small degree of firing rate distortion is acceptable, especially when dither widths do not exceed the temporal scale at which predictable non-stationarities may occur. Others, though, have also argued that dithering may have unintentional consequences on the autostructure of a spike train, thereby rendering its effects on intrinsic ISI distributions and those on coincidence counts indistinguishable (Pipa, Grün, and Vreeswijk, 2013). However, it is not immediately clear that these two characteristics of neuronal firing occur as a result of biologically separable mechanisms. That is, a cell’s ISI distribution is likely tied to the pattern of activity of its inputs, not just to intrinsic properties. Further, these intrinsic rate parameters - such as membrane electrical properties, axon length, and cytoplasm resistance - are likely most significant at the millisecond scale (Kuriscak et al., 2012). Therefore, we observe that distortion of the ISI profile may be a necessary and even desirable consequence of dithering for the purpose of creating a coincidence null distribution that assumes the independence of two spike trains, and that its effects on precise autostructure are temporally distinct. For these reasons, in our view, dithering offers an appropriate and conservative null for synchrony analyses.

### Simulation Procedures

Assembly detection processes are often tested using unique simulation procedures. Here, we have chosen to reflect firing rate diversity by simulating neurons with generally different rates. The participation of these neurons in assemblies was also engineered to reflect their mean rate. In many cases (Berger et al., 2010; Lopes-dos-Santos, Ribeiro, and Tort, 2013; Pipa, Grün, and Vreeswijk, 2013), though, neurons are simulated at 10-20Hz, which exceeds the rate of most principal cells throughout the brain, particularly the cortex (Roxin et al., 2011; Osada, Nishihara, and F. Kimura, 1991). It must likewise be observed that simulating neurons with higher rates can satisfy methodologies with intrinsic upper limits on spike train sparsity, a limitation that we have explicitly attempted to address in Methods. Moreover, when neurons are simulated homogeneously, the question of rate bias in assembly detection methodology may also not be addressed. Throughout all simulations presented herein, we have not noticed any such bias in pairwise synchrony analysis nor assembly detection. Despite these advantages, the simulation procedures described here do not reproduce non-stationarities. While we do not expect our Methods to be significantly biased by heterogeneous rate profiles, further experiments addressing assembly detection in the context of neuronal coupling to cortical oscillations should be performed.

Simulating assemblies presents an additional challenge to that of reflecting baseline neural activity. We have generally followed the assembly simulation procedures used by Berger, observing that they provide ample opportunity to both modulate assembly strength and scalability to accommodate mean firing rates (Berger et al., 2010). The verisimilitude of assembly simulation, though, is largely determined by one’s stated assembly definition. Simulating assemblies as a hidden process with a vector of copy probabilities, for example, reflects the implied stochastic nature of assembly activity. Similarly, assembly activation rates (2Hz) were chosen to limit the contribution of the assembly to the overall spike train. For this reason also, we chose to modulate assembly strength through copy probability, not rate.

### Assembly Detection Algorithms

The measure of pairwise coactivity described herein is a version of CSF proposed by Berger. In that study, Berger et al. examine the possibility of using CSF to limit the number of cells being considered for assembly detection, with the goal of reducing the computational complexity of assigning cells to assemblies (Berger et al., 2010). In a similar vein, we combine CSFs with the coincidence distribution following dithering to detect excess synchrony. Many alternative assembly detection methods, though, rely on dimensionality reduction (e.g. PCA, FA, and ICA) of the spike train matrix to detect assemblies, not a matrix of pairwise measurements (Yu et al., 2009; Peyrache et al., 2009; Lopes-dos-Santos, Ribeiro, and Tort, 2013). We must note that while a pairwise-based approach is not designed to detect higher-order relationships, it would only be expected to fail should these relationships exist in the absence of detectable pairwise synchrony.

Another important aspect of assembly detection, which dimensionality reduction addresses, is that of computational complexity. While our measure and analysis of pairwise synchrony is reasonably efficient in this regard, the assembly detection algorithm would likely struggle to scale efficiently with the size of the recorded population. Several layers of optimization might be implemented to constrain the severity of this problem, with an analytical estimate for the null distribution of *ε* being key in this regard. Nevertheless, the iterative nature of the algorithm would prevent scaling to populations numbering many hundreds of highly connected cells without further optimization, in which the likelihood of discovering many assemblies is expected to increase. To this end, we observe that scalable solutions do not need to forego reliance on pairwise measurements. Graph-based clustering algorithms are particularly well suited in this regard (Billeh et al., 2014; Mölter, Avitan, and Goodhill, 2018; Tavoni, Cocco, and Monasson, 2016), though many may not detect overlapping clusters or would require the number of assemblies to be known in advance (Singh and Garg, 2022; Liu and Barahona, 2020).

### Fine Temporal Motifs

In these experiments we have aimed to characterize temporal spike patterns either with (Fig. 4) or without (Fig. 3) specific attention to conserved order. The existence of temporal motifs is striking but not unexpected from synchronous neurons (R. Kimura et al., 2016; Izhikevich, 2004). As recordings were made from multiple cortical laminae, it is impossible to verify whether the degree of interconnectivity between cells reflects expected inter- and intra-laminar dynamics, nor the impact that common thalamic or cortical afferent pathways may have on determining these motifs. Further, it is noteworthy that about three-quarters of all synchronous pairs of neurons were not active in any particular order. It is possible that this is merely a consequence of small effect sizes: indeed, cross-correlograms for coordinated and uncoordinated synchronous pairs appear similar (Fig. 4F).

Despite these limitations, the proportion of coordinated pairs and evidence of synchrony are indicative of a sparse network and hence assembly separability.

### Encoding Complex Sounds

While suggestions of robust temporal codes have appeared in data from single neurons (Schnupp et al., 2006), other work has found that single units rarely match psychometric performance (Wang et al., 2007; Bizley et al., 2010). Thus, our analysis of small synchrony-driven assemblies suggests that, despite receptive field and response diversity, it is possible to cluster neurons such that trial-by-trial coherence is retained. If synchrony were irrelevant, random assemblies should encode as well as SDAs. Instead, we find a significant advantage for SDAs. From both rate-coding and temporal-coding perspectives, this observation is key in resolving the limitations of single neuron codes without sacrificing sparsity as a general coding principle.

Our analysis of two types of rate codes suggests that synchrony-driven assemblies can retain most of their information in the form of coincidence counts rather than total spike counts. Inevitably, coincidence-based rate codes draw information from fewer spikes, and may thus be an energetically favorable coding mechanism. Synchronous spikes have likewise been shown to carry more information than asynchronous spikes in cat auditory cortex (C. A. Atencio and C. E. Schreiner, 2013; C. Atencio and C. Schreiner, 2016). Moreover, STDP is a known mechanism for extracting information from coincident events, potentially allowing the activity of synchrony-driven assemblies to be easily read out by downstream circuits. It is also interesting to note that temporal codes for complex sounds could retain this mechanism to identify periods of increased assembly activity. Hence, from the decoder’s perspective, SDAs may provide an information-rich low-cost alternative to more densely distributed rate codes.

How neurons could encode the relative timing of assembly activity remains an open question. In the few sessions when multiple fully distinct assemblies were identified, we observed that assemblies could ‘tile’ their responses and exhibit different response patterns. Thus, the ‘gaps’ in a single assembly’s temporal code may be filled by other assemblies acting in parallel (Hahnloser, Kozhevnikov, and Fee, 2002; Buzsáki, 2010). We propose that, rather than encode the times at which a single assembly is active, it may be more feasible to encode short sequences of assembly activity (activity in assembly A immediately followed by B, for example). Therefore, provided that sufficient assemblies exist to encode each aspect of short sounds, a temporal code for complex sounds could be condensed for more efficient downstream computation.

Finally, the significance of temporal codes in random assemblies must also be addressed. Here, we note that temporal information is likely distributed across the population. The coherence of this information is specifically enhanced in synchrony-driven assemblies, leading to an increase in decoder performance upon inclusion of synchronous units. Presumably, there exist other neurons in the population that could form assemblies with remaining cells (those that we have included in random assemblies). The distinction between units included and excluded from our assemblies is thus likely born of the limited number of neurons recorded on each session, not cortical coding strategies.

### Primary and Non-Primary Assemblies

We have observed, throughout this analysis, that A1 and PEG dynamics do not differ significantly in the metrics we have tested. This is broadly consistent with a recent report comparing the responses of primary and non-primary ferret auditory cortices to either synthetic or natural complex sounds, including ferret vocalizations (Landemard et al., 2021). Other results, however, are indicative of significant differences. In one such study, non-primary areas were found to exhibit stimulus habituation that is not present in primary cortex (Lu et al., 2018). In another, non-primary cortex responses were modulated by sound category and engagement in a challenging discrimination task (Heller, Hamersky, and Stephen V David, 2024). Together, these results indicate that the separability between primary and non-primary responses may be enhanced by specific task types, and possibly by task difficulty. When passive listening is sufficient, responses in the two regions are mostly similar. Thus, to determine whether the coding strategies we have investigated truly characterize both cortices, an important next step int he form of task-based analysis will be required.

## Conclusions

The graph-based assembly detection method we describe was shown to depend on a reliable analysis of precise pairwise synchrony, to be robust to jitter, to tolerate overlapping communities, and to work efficiently in sparse spike matrices built from heterogeneous low-to-mid-rate cells. This method is by no means the first to organize cells as a graph (R. Kimura et al., 2016; Billeh et al., 2014; Tavoni, Cocco, and Monasson, 2016; Mölter, Avitan, and Goodhill, 2018), and the properties of our chosen surrogate data generation procedure for pairwise synchrony are likewise well documented in other work (Pipa, Wheeler, et al., 2008; Louis, 2010; Pipa, Grün, and Vreeswijk, 2013). To the best of our knowledge, however, this analytical null distribution of independent two-spike dithering following a uniform discrete random variable is not presented elsewhere, nor is this approach used for assembly analysis. By combining this statistical framework with graph-based assembly detection, we show that identified assemblies exhibit stereotypical temporal dynamics and reliably encode complex sounds. Further, we have shown that these assemblies exhibit stereotypical temporal dynamics and reliably encode complex sounds, likely through a combination of coincidence-based rate codes and temporal codes. These observations establish a foundation for studying assemblies at the temporal resolution required to infer interconnectivity, and ultimately to probe the cognitive computations that such coordinated networks may support. Looking forward, assembly analysis will be key to understanding how cortical networks overcome biophysical noise to generate reliable representations of complex, naturalistic stimuli.

## Appendix A

By summing all weights, we sum all values that Eq.13 may take for all recorded neurons. Observe that:

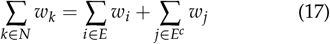

where *N* is the set of all recorded neurons. Also:

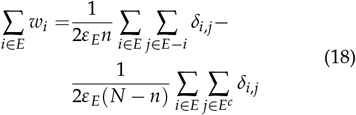

from Eq.13. Similarly, for all cells not in the assembly:

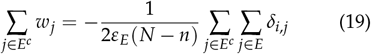

Crucially, the first double sum in Eq.18 represents 2 ∑ *I*, while the second, as well as that in Eq.19, are ∑ *O*. Integrating this information into Eq.17, we obtain:

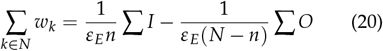

Factoring 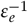, we have:

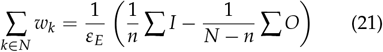

Observing that the term in parentheses is indeed our definition of *ε*_*E*_ proves the proposition.

## Supplementary

**Figure S1:**
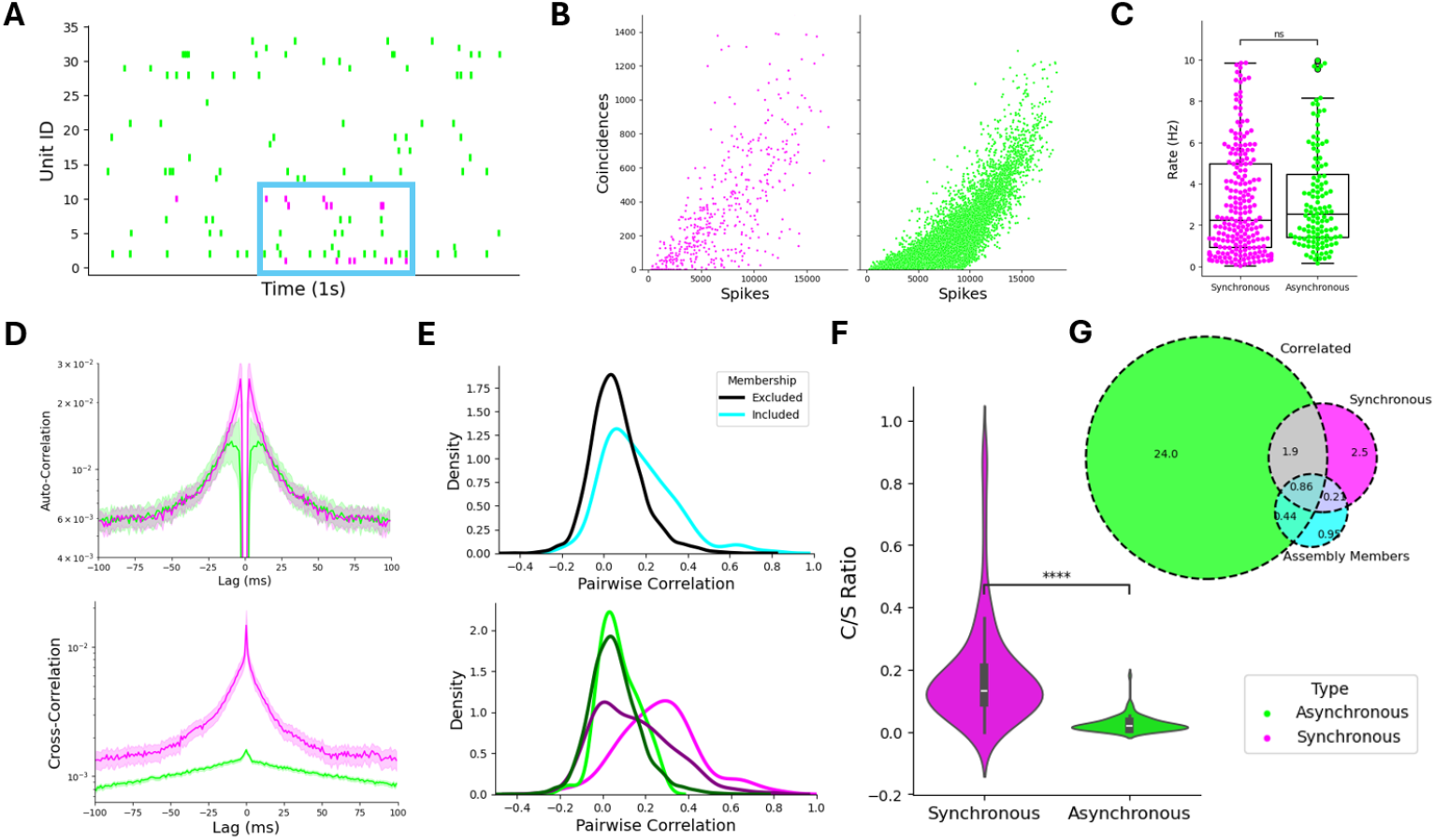
Adirectional pairwise temporal dynamics in synchronous and asynchronous units. A) Example raster plot from PEG recording session, with one four-unit assembly shown in purple. The assembly appears to be synchronously active several times in this window, with all or most units participating (cyan box). B) Coincidence counts and spike counts for synchronous (purple) and asynchronous (green) pairs. Observe that, on average, synchronous pairs generate more coincident events per spike. C) Distribution of mean firing rates for neurons synchronous with at least one other unit (left) and those that are not synchronous with any other cell (right). D) Auto-correlogram (top) and cross-correlogram (bottom) for synchronous (purple) and asynchronous (green) units. Synchronous units are defined as in C for the auto-correlogram. E) Distribution of trial-by-trial pairwise correlation between pairs included in detected assemblies (cyan) and those that remain excluded (black). The bottom panel further differentiates between synchronous and asynchronous pairs (dark for excluded, light for included). F) Distribution of coincidence counts per spike (C/S) ratio, quantifying the pairwise relationship shown in D. G) Venn-diagram showing pairs of neurons that are correlated (green), synchronous (purple), and included in assemblies (cyan). Numbers indicate the % of all pairs that fall into a given category.

**Figure S2:**
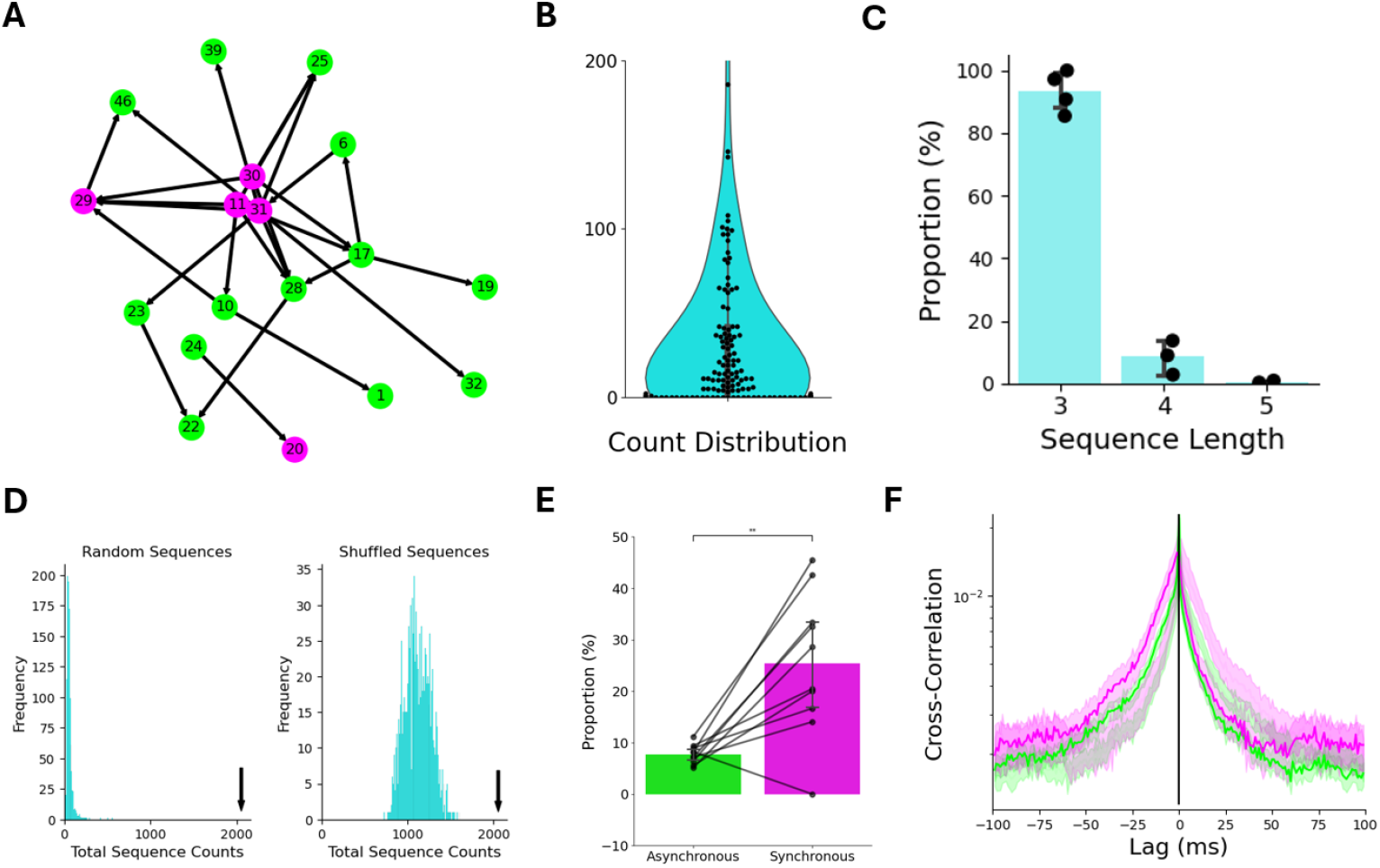
Pairwise temporal dynamics in synchronous and asynchronous PEG units. A) Network representation of an example PEG session, with arrows connecting pairs of units that are both synchronous and temporally coordinated (from pre to post). Assembly members are shown in purple. B) Frequencies of predicted sequences, with predictions made by following ‘paths’ on graphs like those in A. Data from all sessions with at least 5 predictions is shown. C) Proportion of identified predicted sequences by size, per session. The overwhelming majority are short (3 units), with 1-10% consisting of four units and the remainder five. D) Example null distributions of total sequence counts for either random sequences (left) of shuffled predicted sequences (right), generated from the same session as A. In both cases, predicted sequence counts are significantly greater than those of surrogates. E) Proportion of asynchronous or synchronous cell pairs that exhibit significant temporal coordination at 5ms intervals (defined by the pre-post binomial test). Data is shown per session. F) Cross-correlogram for ordered coordinated (purple) and non-coordinated (green) synchronous pairs, for all sessions. The mirror image of each correlogram is plotted as a ‘shadow’ reflected over the black line to facilitate visual comparisons.

**Figure S3:**
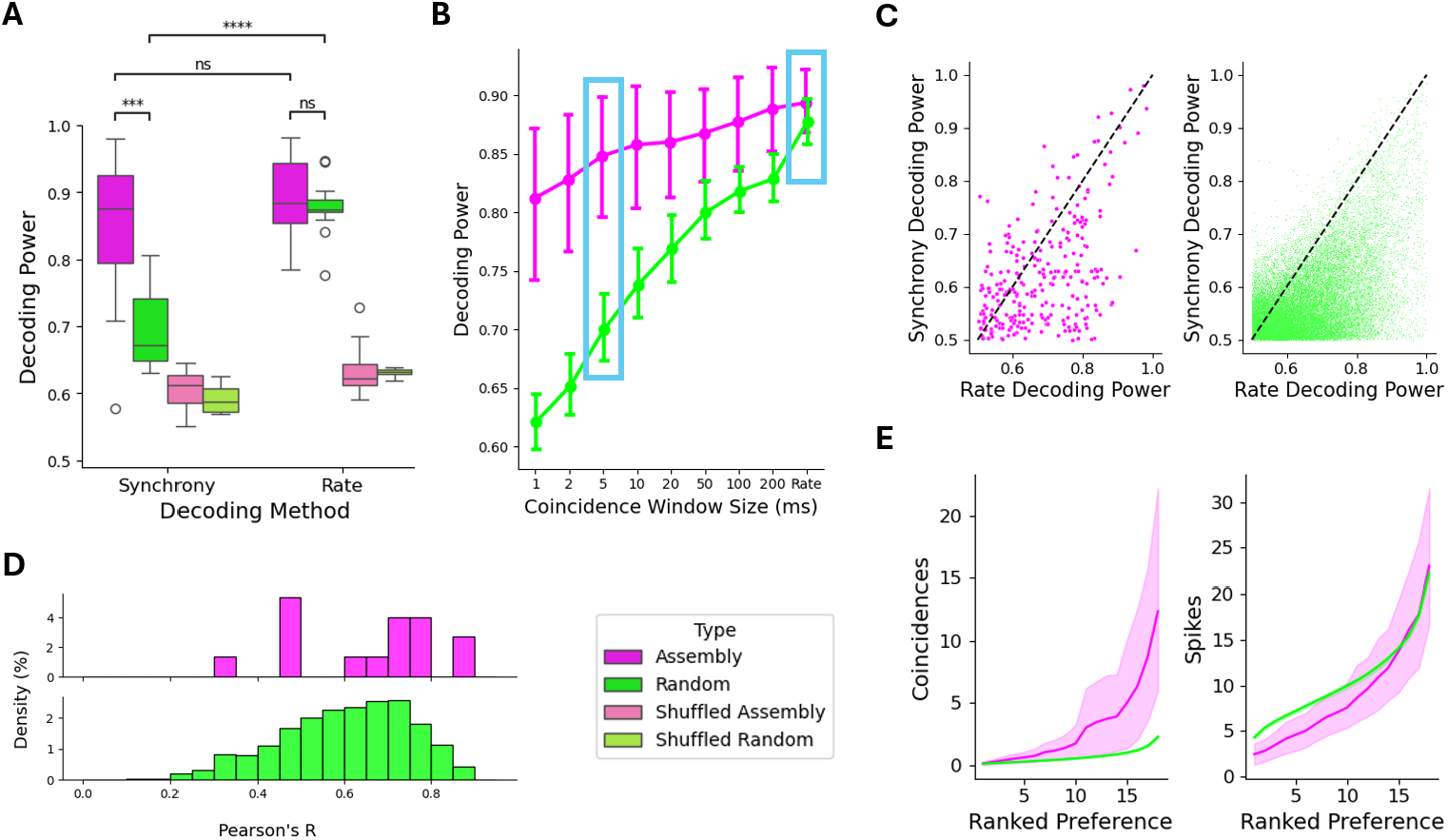
Synchrony-driven assemblies preserve information from rate codes on millisecond timescales in PEG. A) Results of ROC decoding analysis, with the area under the curve (AUC) plotted as decoding power for the best decoded stimulus per assembly. Decoding from assembly (n = 15) responses is shown in purple, with surrogates plotted in green. 100 random surrogates were generated per identified assembly. Pale colors show results from trial-shuffling. B) Mean decoding power (AUC) for preferred stimulus with varying maximum interval. Cyan boxes indicate ‘Synchrony’ and ‘Rate’ data shown in panel A. C) Comparison between spike count (rate) and coincidence count (synchrony) decoding for assemblies (left) and random surrogates (right). Dashed lines indicate the first quadrant bisector. Data from all decoded stimuli is shown. D) Distribution of Pearson’s correlation between coincidence count and spike count responses. E) Mean number of coincidences (left) or spikes (right) per stimulus, ordered by response strength per assembly.

**Figure S4:**
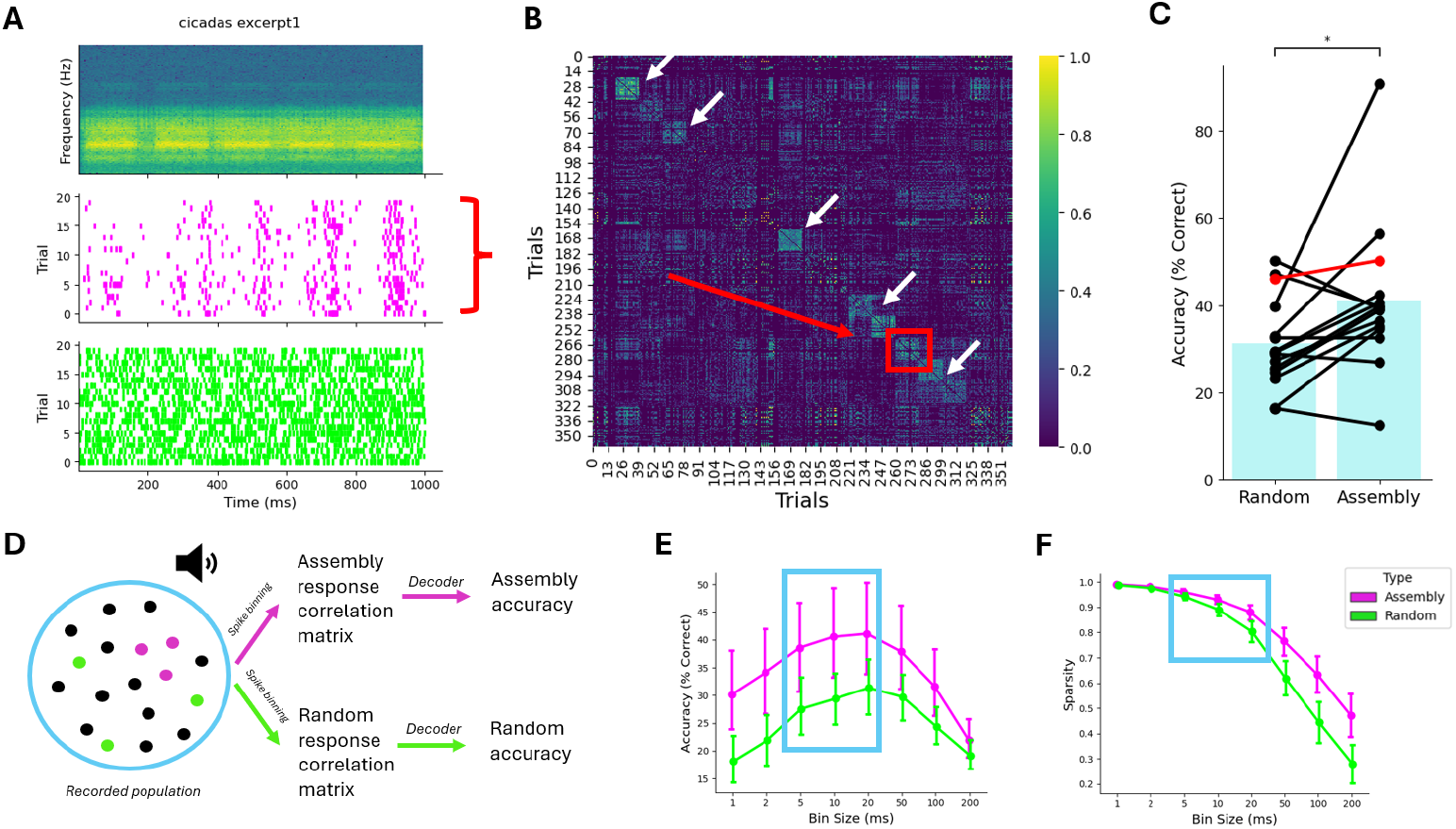
PEG synchrony-driven assemblies employ sparse temporal codes. A) An example stimulus spectrogram (top) with associated trial-by-trial raster plots for one assembly (middle) and all other units (bottom). B) Correlation between assembly response vectors, binned at 20ms, for all repetition trials from one recording session. The example in A is marked in red. Note that trial order is not chronological, and has been rearranged to highlight consistent responses (white arrows). C) Temporal decoding accuracy for assembly-surrogate pairs (n = 15). The example from A is marked in red. D) Diagram of temporal decoding, with assembly and surrogate spike trains being used to generate a correlation matrix that is then decoded to determine accuracy (see Methods). E) Temporal decoding mean accuracy with varying bin width. F) Sparsity - defined as the proportion of empty bins - for assemblies (purple) and surrogates (green) with varying bin width.

**Figure S5:**
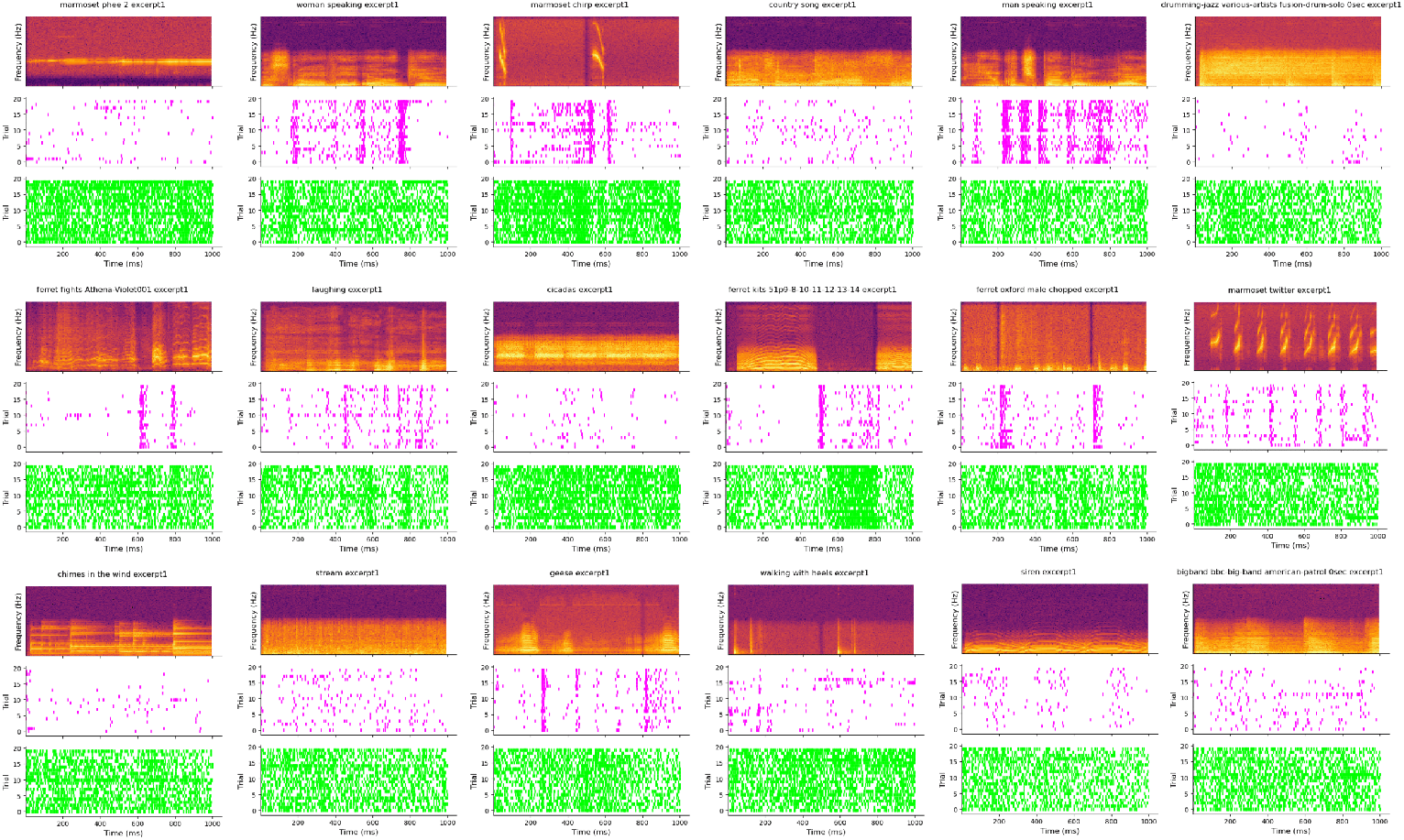
Stimulus spectrograms (top) with associated trial-by-trial raster plots for one assembly (middle) and all other units (bottom). All repeat stimuli from one A1 recording session are shown. Notice the robust temporal patterns in roughly half of all panels.

**Figure S6:**
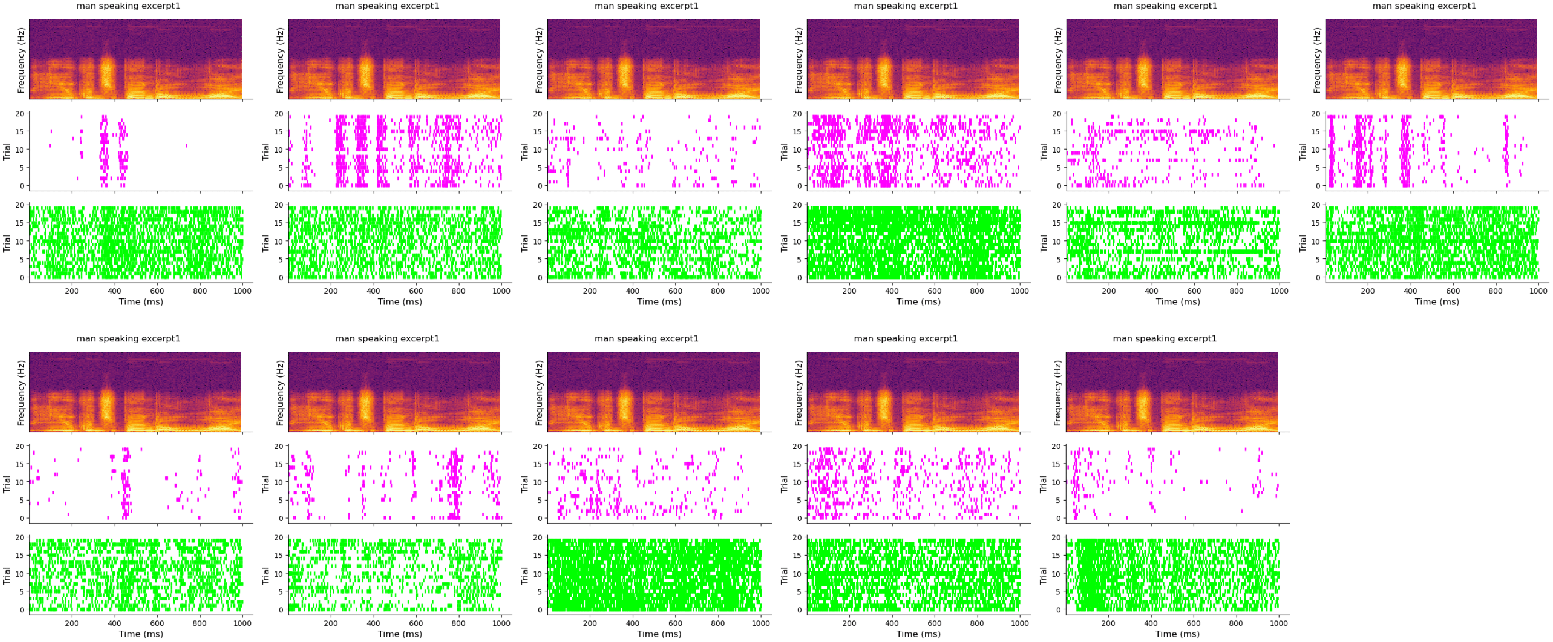
Stimulus spectrograms (top) with associated trial-by-trial raster plots for one assembly (middle) and all other units (bottom). Here, we show all assembly responses across all A1 recording sessions to one example stimulus. Notice the variable temporal patterns produced by different assemblies.

